# Genomic Characterization and Therapeutic Potential of the Lytic Bacteriophage Curly against *Klebsiella pneumoniae* in Human Innate Immune Cells and a Murine Pneumonia Model

**DOI:** 10.64898/2026.08.25.747058

**Authors:** Mounika Duggineni, Sitaramaraju Adduri, Rajesh Mani, Leticia Guzman Ruiz, Andy Omeje, Narendra Kumar Gonepudi, Joshua K. Kleam, Mounika Kumaraswamy, John J. Dennehy, Guohua Yi

## Abstract

*Klebsiella pneumoniae* is an important cause of severe respiratory and systemic infections, and the increasing prevalence of multidrug-resistant strains has created an urgent need for alternative antibacterial strategies. In this study, nine *K. pneumoniae*-infecting bacteriophages isolated from diverse environmental sources were characterized genomically and functionally. Genome analyses revealed substantial genomic and proteomic diversity among the isolates. Functional screening against the clinical *K. pneumoniae* isolate JJD85 identified Curly as the most active phage, producing the highest plaque-forming titer and rapid suppression of bacterial growth in liquid culture. Curly was predicted to have a virulent lifestyle and encoded structural, genome-packaging, and DNA replication-associated proteins. In primary human monocyte-derived macrophage cultures, Curly markedly reduced bacterial burden in both cell-associated and cell-free fractions, while treatment of primary human neutrophil cultures produced an approximately 10^6-fold reduction in total recoverable bacterial burden. Transmission electron microscopy demonstrated phage-like particles within bacterial profiles located in both extracellular and macrophage-associated intracellular compartments. In a C57BL/6J murine pneumonia model, intranasal Curly treatment reduced pulmonary bacterial burden in a dose-associated manner, with approximately 10-fold and 100-fold reductions at the low and high doses, respectively. Curly treatment also attenuated infection-associated lung inflammation and preserved pulmonary architecture. These findings identify Curly as a promising bacteriophage candidate against *K. pneumoniae* and support further evaluation of its host range, resistance profile, and therapeutic potential.

## Introduction

Klebsiella pneumoniae is an encapsulated Gram-negative bacterial pathogen that causes a broad spectrum of infections, including pneumonia, urinary tract infection, bacteremia, meningitis, and liver abscess^1,2^. Severe infections occur particularly in hospitalized and immunocompromised individuals and are increasingly difficult to treat because of the emergence and dissemination of multidrug-resistant strains^1,2^. The ability of K. pneumoniae to acquire antimicrobial resistance determinants through mutation and horizontal gene transfer has contributed substantially to the global antimicrobial resistance burden^3,4^. Of particular concern are extended-spectrum β-lactamase-producing^5,6^ and carbapenem-resistant K. pneumoniae^7,8^, for which therapeutic options can be severely limited. These challenges have intensified the search for alternative antibacterial approaches capable of targeting drug-resistant K. pneumoniae^3^.

Bacteriophages are viruses that specifically infect bacteria and have re-emerged as potential therapeutic agents for difficult-to-treat bacterial infections^9–11^. Virulent, or lytic, phages replicate within susceptible bacterial hosts and ultimately induce bacterial lysis, providing a mechanism of bacterial killing that is distinct from conventional antibiotics^12–14^. Their capacity for host-specific killing, replication in the presence of susceptible bacteria, and activity against antibiotic-resistant organisms make them attractive candidates for treatment of multidrug-resistant infections^10,11,13^. Preclinical studies have demonstrated therapeutic activity of phages against *K. pneumoniae* in several infection settings, including experimental pneumonia^15,16^. Nevertheless, successful translation of phage therapy remains constrained by variability in host range, emergence of phage resistance, differences in phage stability and tissue accessibility, and the influence of dose, route of administration, and host immune responses on treatment efficacy^17^.

The extensive genetic and phenotypic diversity of both *K. pneumoniae* and its bacteriophages represents an additional challenge for therapeutic development^18,19^. Individual phages commonly infect only subsets of bacterial strains, while even related phages can differ considerably in host recognition and antibacterial activity^19,20^. Consequently, establishing genetically diverse and well-characterized phage collections is important both for identifying highly active individual candidates and for the future development of rational phage cocktails with complementary host ranges^19,21^.

Genomic characterization is particularly important for defining relationships among candidate phages, identifying structural and replication-associated genes, and screening for genomic features that may be undesirable for therapeutic use^10,21^. However, genomic relatedness alone cannot establish therapeutic potency, emphasizing the need to combine comparative genomics with direct functional screening^20,22^.

A further consideration is that antibacterial activity measured in conventional bacterial cultures may not fully predict phage performance in the infected host^11,23^. During pulmonary *K. pneumoniae* infection, bacteria encounter macrophages and neutrophils, two major components of innate immune defense^24,25^. Macrophages participate in early recognition and phagocytic clearance of bacteria, whereas neutrophils are rapidly recruited and contribute to bacterial killing through phagocytosis and additional antimicrobial mechanisms^24,25^. *K. pneumoniae*, however, possesses multiple mechanisms that enable resistance to innate immune clearance and persistence within the infected host^24,26^.

Thus, determining whether candidate phages retain antibacterial activity in the presence of primary human immune cells provides an important intermediate assessment between bacterial culture assays and animal models^23^.

In the present study, we characterized a panel of K. pneumoniae-infecting bacteriophages isolated from diverse environmental sources using whole-genome nucleotide and proteomic analyses and compared their antibacterial activities against the clinical K. pneumoniae isolate JJD85. From this collection, we identified Curly as a lead phage based on its strong plaque-forming and bactericidal activity and further characterized its predicted genomic architecture and lytic lifestyle. We then evaluated Curly in progressively more biologically relevant experimental systems, including primary human monocyte-derived macrophages and neutrophils. Transmission electron microscopy was used to examine the spatial relationship among macrophages, bacteria, and phage particles, and therapeutic activity was subsequently evaluated in a C57BL/6J mouse model of K. pneumoniae pneumonia. This integrated approach was designed to determine whether a phage selected through genomic and functional screening retains antibacterial activity across bacterial culture, primary human immune-cell, and in vivo pulmonary infection settings.

## Methods

### 1. Bacterial and bacteriophage isolates

*Klebsiella pneumoniae* strain JJD85, isolated from hospital wastewater, was used as the bacterial host throughout this study. Nine *K. pneumoniae*-infecting bacteriophages isolated from diverse environmental and animal-associated sources were evaluated. Chickie was isolated from chicken coop soil, Piggy and Moe from pig feces, Sheepy from sheep pen soil, Curly from horse feces, and Vulcan from turkey vulture feces. Fei, Reina, and Malika were isolated from hospital wastewater.

### Genome sequence and comparative genomic analysis

Complete genome sequences of the nine bacteriophage isolates were subjected to comparative genomic analysis. Pairwise whole-genome nucleotide similarities were calculated using the Virus Intergenomic Distance Calculator (VIRIDIC). The resulting intergenomic similarity matrix was visualized as a heatmap, and hierarchical clustering was performed using the complete-linkage clustering method. Proteome-based relationships among the phages were further evaluated using ViPTree v4.1. Analyses were performed using the dsDNA virus dataset with prokaryotes selected as the host type. Protein-coding genes were predicted using Prodigal with genetic code corresponding to the Bacterial, Archaeal and Plant Plastid Code. Proteomic relationships between the study phages and reference viruses were assessed using the ViPTree-generated proteomic tree, and pairwise genome comparisons were examined using the corresponding translated sequence similarity alignments. PhageAI was used to predict phage lifestyle.

### Amplification and quantification of bacteriophages

Bacterial culture and bacteriophage propagation were performed following the protocols described by Bonilla et al^27^. K. pneumoniae strains were cultured overnight in Luria–Bertani (LB) broth at 37 °C with shaking at 250 rpm. To propagate bacteriophages,5 × 10¹⁰ PFU/mL of phages were mixed with 100 μL of K. pneumoniae (5 × 10⁸ CFU/mL) and inoculated into LB broth. This mixture was incubated at 37 °C with agitation for approximately 4 h until complete lysis was observed, as evidenced by clearing of the culture. The resulting lysate was collected and centrifuged at 4,000 × g for 20 min to remove bacterial debris. The supernatant was carefully transferred to a new sterile tube and filter-sterilized using a 0.22 μm filter to obtain a cell-free bacteriophage lysate, which was stored at 4 °C until further uses.

For concentration and purification, the bacteriophage lysates were loaded into an Amicon ultrafiltration device 100 kDa and centrifuged at 4,000 × g in multiple rounds until the total volume was reduced to <10 mL. The concentrate was washed twice with bacteriophage buffer (10 mM Tris, pH 7.5; 10 mM MgSO₄; 0.4% NaCl) under the same centrifugation conditions to remove residual contaminants. The final concentrated and purified bacteriophage lysate was recovered, and bacteriophage titers were determined by plaque assay and recorded as PFU/mL.

To determine bacteriophage titers, K. pneumoniae was used as the bacterial host in standard double-layer agar plaque assays. Briefly, 7 mL of LB top agar (3 parts LB broth + 1 part of 0.8% Top agar) was prepared and maintained in a water bath at 45–47 °C to prevent solidification. Bacteriophage samples were serially diluted by adding 100 μL of bacteriophage lysate to 900 μL of bacteriophage buffer, with thorough mixing each step. The diluted bacteriophage suspensions were incubated at 37 °C for 5 min. Subsequently, 100 μL of mid-log phase K. pneumoniae culture (OD600 is 0.5-0.6) was added to the molten top agar, mixed gently by swirling, and poured evenly onto LB agar plates to form a uniform bacterial lawn. After the top agar solidified at room temperature (∼10 mins), 100 μL aliquots of each serially diluted bacteriophage were applied onto the surface of the plates and allowed to absorb completely. The plates were then invertedly incubated overnight at 37 °C, after which visible plaques were counted and used to calculate bacteriophage titers, expressed as plaque-forming units per milliliter (PFU/mL).

Infection assay to determine bacteriophage infectivity: For infection assays, K. pneumoniae was grown to mid-log phase and adjusted to a single-cell suspension of approximately 5*10⁸ cells/mL. These cells were infected with bacteriophages at a multiplicity of infection (MOI) of 1 (one PFU of bacteriophage per bacterial cell) and incubated at 37°C for 10 minutes. Following adsorption, the infection mixture was added to 7 mL of 0.8% top agar and spread onto 150-mm LB agar plates for overnight incubation at 37°C. The presence of plaques on the bacterial lawn confirmed successful bacteriophage infection and lytic activity. Bacteriophage titers in the original lysates or post-infection samples were determined using standard plaque assay as described above.

### In Vitro lytic activity of bacteriophages against K. pneumoniae in liquid culture

The lytic activity and infectivity of bacteriophages against K. pneumoniae was evaluated in liquid cultures by monitoring bacterial growth kinetics overtime via optical density at 600 nm (OD600). K. pneumoniae cultures were grown in LB broth at 37 °C with shaking to mid-exponential phase (OD₆₀₀ is approximately 0.5–0.6). For infection, 10 μL of K. pneumoniae (5 × 10⁸ CFU/mL) was mixed with 10 μL of high-titer bacteriophage lysate (5 × 10¹⁰ PFU/mL). This yielded an approximate multiplicity of infection (MOI) of 100 calculated as the number of bacteriophage particles per bacterial cell. We used a high MOI to ensure simultaneous infection of most bacterial cells, facilitating rapid and detectable lysis. The mixture was incubated under standard growth conditions, and bacterial growth was monitored over time by measuring OD₆₀₀ at each hour for five more hours.

### Human samples

The study was approved by the Institutional Review Board of The University of Texas Health Science Center at Tyler (IRB #2024-072). Peripheral blood was collected from four healthy adult donors (n = 4), and written informed consent was obtained from all participants before sample collection. All procedures involving human samples were performed in accordance with the approved IRB protocol.

### Flow cytometry

Macrophages and neutrophils were characterized using flow cytometry based on surface and intracellular marker expression. Single-cell suspensions were prepared and maintained on ice throughout the staining procedure. Approximately 2 × 10⁵ to 1 × 10⁶ cells were aliquoted per sample, centrifuged at 450 × g for 10 min, and then the supernatant was removed. To reduce non-specific antibody binding, Fc receptors were blocked using Human TruStain FcX (BioLegend) diluted 5:100 in Phosphate Buffered Saline (PBS) and incubated for 20–30 min at 4°C. The cells were then washed with PBS and centrifuged at 450 × g for 10 min. Cell viability was assessed using a fixable Live/Dead Near-IR dye (BioLegend), diluted 1:100 in FACS buffer (PBS containing 2% FBS), and incubated for 20–30 min at 4°C in the dark. Following incubation, the cells were washed and centrifuged. Surface staining was performed by incubating cells with fluorochrome-conjugated antibodies diluted 1:100 in FACS buffer for 20–30 min at 4°C. Macrophages were identified using Brilliant Violet 480 CD14 (eBioscience Cat # 414-0149-42) and neutrophils were identified using a APC CD66⁺ marker (Biolegend cat # 305117). Cells were fixed using True-Nuclear™ Fix Concentrate (1:4 dilution in Fix Diluent) and incubated for 30–45 min at room temperature in the dark. Finally, the cells were centrifuged at 450 × g for 5 min, and then the supernatant was removed. All samples were resuspended in 400 µL of flow cytometry staining buffer and acquired on the flow cytometer Attune NxT (Acoustic Focusing Cytometer) using the excitation lasers Blue (488 nm) and Red (638 nm), and the emission filters Blue 574/26 nm and Red 720/30 nm for CD14 and CD66, respectively.

Appropriate compensation controls were obtained. Data were analysed using standard gating strategies: initial gating on forward and side scatter to exclude debris, followed by singlet discrimination and live/dead exclusion. Macrophages were identified as CD14⁺ viable cells, while neutrophils were identified as CD66⁺ viable cells.

### Evaluating bacteriophages-mediated killing of K. pneumoniae in macrophages

**1) Isolation of CD14+ monocytes from human peripheral blood:** Bacteriophage activity was evaluated as described previously [47]. Peripheral blood was collected from four healthy donors into sodium heparin tubes. The study was approved by Institutional Review Board (IRB) of University of Texas at Tyler Health science Center. Informed consent was obtained from all participants following Helsinki declaration 1975. Peripheral blood mononuclear cells (PBMCs) were isolated by Ficoll-Paque Plus (17-1440-02, GE Healthcare, Danderyd, Sweden) density gradient centrifugation at 400 × g for 30 min at room temperature. The PBMC layer was collected and washed twice with PBS by centrifugation at 400 × g, 4 °C for 10 min. CD14⁺ monocytes were then isolated from PBMCs by negative immunomagnetic selection using human CD14⁺ microbeads (EasySep™ Human Monocyte Enrichment Kit without CD16 Depletion, Catalog #19058) in PBS containing 2% FBS and 1 mM EDTA, following the manufacturer’s instructions. After isolation, CD14⁺ monocytes were washed with 1 mL of cold PBS, and cell purity was assessed by flow cytometry (typically >90% CD14+).
**2) Differentiation of CD14+ monocytes to macrophages:** For differentiation, 0.5 × 10⁶ CD14⁺ monocytes were seeded into each well of a 24-well plate containing 1.5 mL of RPMI-1640 medium supplemented with 10% FBS and 1% penicillin–streptomycin. Cells were cultured at 37 °C in 5% CO₂ in the presence of 50 ng/mL GM-CSF (02532, Stem Cells, Vancouver, Canada) and 50 ng/mL M-CSF (216-MC, R&D Systems, Minneapolis, USA). Cytokines were replenished once on day 3. On day 6, a subset of cells was collected and stained for macrophage markers to confirm differentiation by flow cytometry. The remaining cells were washed with PBS and rested overnight in fresh medium without serum or antibiotics to minimize background interference in subsequent infection assays.
**3) Macrophage infection and bacteriophage treatment:** For infection assays, the differentiated macrophages were infected with K. pneumoniae at a multiplicity of infection (MOI) of 5 (bacteria-to-cell ratio). One-hour post-infection, the supernatant and adherent macrophages were collected to quantify bacteria in cell free fraction and cell associated fraction, respectively, and the wells without bacteriophage addition were used as controls. Subsequently, bacteriophage #418 was added at an MOI of 10 to designated wells, while control wells received no bacteriophage. The plates were incubated at 37 °C. Bacterial growth, both cell free fraction and cell associated fraction, was monitored at 1 h and 4 h post-bacteriophage addition.

To quantify cell associated fraction, culture medium was removed, and cells were washed with PBS to remove all cell free fraction bacteria. Macrophages were then lysed by incubation with ddH₂O for 10 min for hypotonic shock, followed by 0.1% Triton X-100 for 10 min to ensure complete disruption of the macrophages. Serial dilutions of the lysates were prepared, and 100 μL of each dilution was plated on LB agar plates. To quantify the bacteria in cell free fraction, supernatants from different time points were serially diluted and plated directly. Plates were incubated overnight at 37 °C, and bacterial colonies were counted to determine colony-forming units (CFU).

### Evaluating bacteriophage killing of K. pneumoniae in neutrophils

**1) Isolation of neutrophils from human peripheral blood:** Neutrophils were isolated from human peripheral blood using the EasySep™ human Neutrophil Isolation Kit (Stemcell Technologies, Cat# 17957) according to the manufacturer’s instructions. Briefly, whole blood was collected from healthy donors in tubes containing anticoagulant. A standard density gradient separation using Ficoll-Paque was performed. The plasma layer, mononuclear cell band, and density gradient medium were carefully removed and discarded, leaving only the erythrocytes/red blood cell (RBC) pellet containg neutrophils intact. Residual RBCs were lysed by adding 9 volumes of ice-cold ammonium chloride lysis solution to one volume of the RBC pellet, mixing thoroughly, and incubating on ice for 15 min. The sample was centrifuged at 500 × g for 10 min with the brake set to low to minimize cell disruption. The supernatant containing lysed RBC debris was discarded, and the cell pellet (enriched with neutrophils) was washed with cold (2–8 °C) RPMI complete medium followed by centrifugation at 120 × g for 10 min with the brake off. After discarding the supernatant, the cell pellet was resuspended, and cell number was determined using the trypan blue exclusion assay with an automated cell counter. The cells were then pelleted again and resuspended at 5 × 10⁷ cells/mL in cold RPMI complete medium, and neutrophil isolation was carried out according to the manufacturer’s instructions. Purity of the isolated neutrophils was confirmed by flow cytometry analysis. Isolated neutrophils were seeded at 5 × 10⁵ cells/well in 12-well plates containing RPMI complete medium and incubated overnight.
**2) In vitro neutrophil infection and bacteriophage treatment:** On the following day, the cells were infected with K. pneumoniae at a multiplicity of infection (MOI) of 5. One-hour post-infection, centrifuged at 450g for 10min and discard the supernatant and the cells were washed twice with PBS to remove extracellular bacteria and then cultured in fresh medium with or without bacteriophage at an MOI of 10 (5 × 10⁷ PFU). After 4 h and 6 h of incubation, both cells and supernatants were collected. The samples were serially diluted from 10⁻¹ to 10⁻⁷ and plated on LB agar plates. Colony-forming units (CFUs) were counted following overnight incubation at 37 °C.

### Transmission electron microscopy

Samples were fixed in a solution containing 4% paraformaldehyde, 2.5% glutaraldehyde, and 0.2% picric acid in 0.1 M phosphate buffer. Following fixation, samples were rinsed with 0.1 M phosphate buffer and post-fixed for 1 h in 1% osmium tetroxide and 1.5% potassium ferricyanide prepared in 0.1 M sodium cacodylate buffer. Samples were then rinsed with water and incubated overnight at 4 °C in an aqueous solution containing 0.5% uranyl acetate and 25% methanol. The following day, samples were rinsed with water and incubated in lead L-aspartate for 30 min at 60 °C. After an additional water rinse, samples were dehydrated through a graded ethanol series consisting of 50%, 70%, 85%, 95% twice, and 100% ethanol four times, with each dehydration step performed for 8 min. Samples were then incubated in propylene oxide for 10 min and infiltrated for 2 h with a 2:1 mixture of propylene oxide and Epon resin. This was followed by overnight infiltration with a 1:2 mixture of propylene oxide and Epon resin under vacuum. Samples were subsequently infiltrated with 100% Epon resin under vacuum twice for 2 h each, embedded in fresh Epon resin, and polymerized at 60 °C for 48 h.

Polymerized blocks were manually trimmed and ultrathin sections were prepared using a diamond knife on a Leica Ultracut 7 ultramicrotome. Sections were collected onto Formvar-coated copper grids for transmission electron microscopy.

### Murine model of *Klebsiella pneumoniae* pulmonary infection and bacteriophage treatment

All animal procedures were approved by the Institutional Animal Care and Use Committee (IACUC) of The University of Texas at Tyler. Wild-type C57BL/6J mice, 6–7 weeks of age, were obtained from The Jackson Laboratory and maintained under standard housing conditions with unrestricted access to food and water. Mice were acclimated for at least 5 days before experimentation and were euthanized if predefined humane endpoint criteria were met. To establish the pulmonary infection model and determine an appropriate bacterial challenge dose, age- and sex-matched mice were anesthetized with isoflurane and inoculated intranasally with either 1 × 10^8^ CFU or 1 × 10^9^ CFU of the clinical *K. pneumoniae* isolate JJD85 suspended in sterile PBS in a total volume of 50 µL. Uninfected control mice received PBS alone. Body weight, clinical signs, and survival were monitored daily. At 2, 4, and 8 days after infection, mice from each infected group were euthanized (n = 4 per dose per time point), and the lungs were aseptically collected. Left lung lobes were homogenized in sterile PBS, serially diluted, and plated on Luria–Bertani (LB) agar. Colonies were enumerated following overnight incubation at 37 °C and bacterial burden was expressed as CFU per lung. Right lung lobes were fixed in 10% neutral-buffered formalin, processed for paraffin embedding, sectioned, and stained with hematoxylin and eosin (H&E) for histopathological examination. Uninfected control mice (n = 4) were processed in parallel at day 8.

For evaluation of therapeutic efficacy, C57BL/6J mice were assigned to four groups (n = 7 per group): uninfected controls, JJD85-infected untreated controls, JJD85-infected mice treated with low-dose Curly, and JJD85-infected mice treated with high-dose Curly. Mice were anesthetized with isoflurane and challenged intranasally with 1 × 10^9^ CFU of JJD85 in 50 µL PBS on two consecutive days. Curly was subsequently administered intranasally once daily for three consecutive days at nominal multiplicities of infection (MOIs) of 1 or 10 for the low- and high-dose treatment groups, respectively. Untreated infected mice received the corresponding vehicle, and uninfected mice served as baseline controls. Body weight, clinical signs, and survival were monitored daily throughout the experiment. At the experimental endpoint, mice were euthanized and lungs were collected aseptically. Lung tissue designated for bacterial quantification was homogenized in sterile PBS, serially diluted, and plated on LB agar, and colonies were enumerated following overnight incubation at 37 °C. The remaining lung tissue was fixed in 10% neutral-buffered formalin, paraffin embedded, sectioned, and stained with H&E for histopathological evaluation.

## Results

### Whole-genome nucleotide comparison reveals distinct genomic groups among the isolated phages

Nine bacteriophages isolated from diverse environmental sources were selected for whole-genome sequencing and comparative genomic analysis. The isolates originated from chicken coop soil, sheep pen soil, horse and pig feces, turkey vulture feces, and hospital wastewater, and their genome sizes ranged from 48,920 to 175,994 bp. All genomes are unique and are not identical to any previously reported isolates. The phages segregated according to genome length, with the smaller-genome phages Curly and Moe clustering together and remaining distinct from the other phages, all of which possessed substantially larger genomes **(Figure 1A)**. Pairwise whole-genome nucleotide similarity was assessed using VIRIDIC **(Figure 1A)**. VIRIDIC analysis revealed marked genomic diversity among the nine phages and identified several distinct groups of closely related isolates. Sheepy, Malika, Piggy, and Vulcan formed a highly related cluster, with pairwise intergenomic similarities ranging from 91.7% to 96.4% **(Figure 1A)**. The greatest similarity within this group was observed between Piggy and Vulcan (96.4%), followed by Malika and Piggy (94.2%), Sheepy and Piggy (93.1%), Malika and Vulcan (92.9%), Sheepy and Malika (92.8%), and Sheepy and Vulcan (91.7%) **(Figure 1A)**. Consistent with this close genomic relationship, all four phages were independently assigned to the genus *Slopekvirus*.

**Figure 1:**
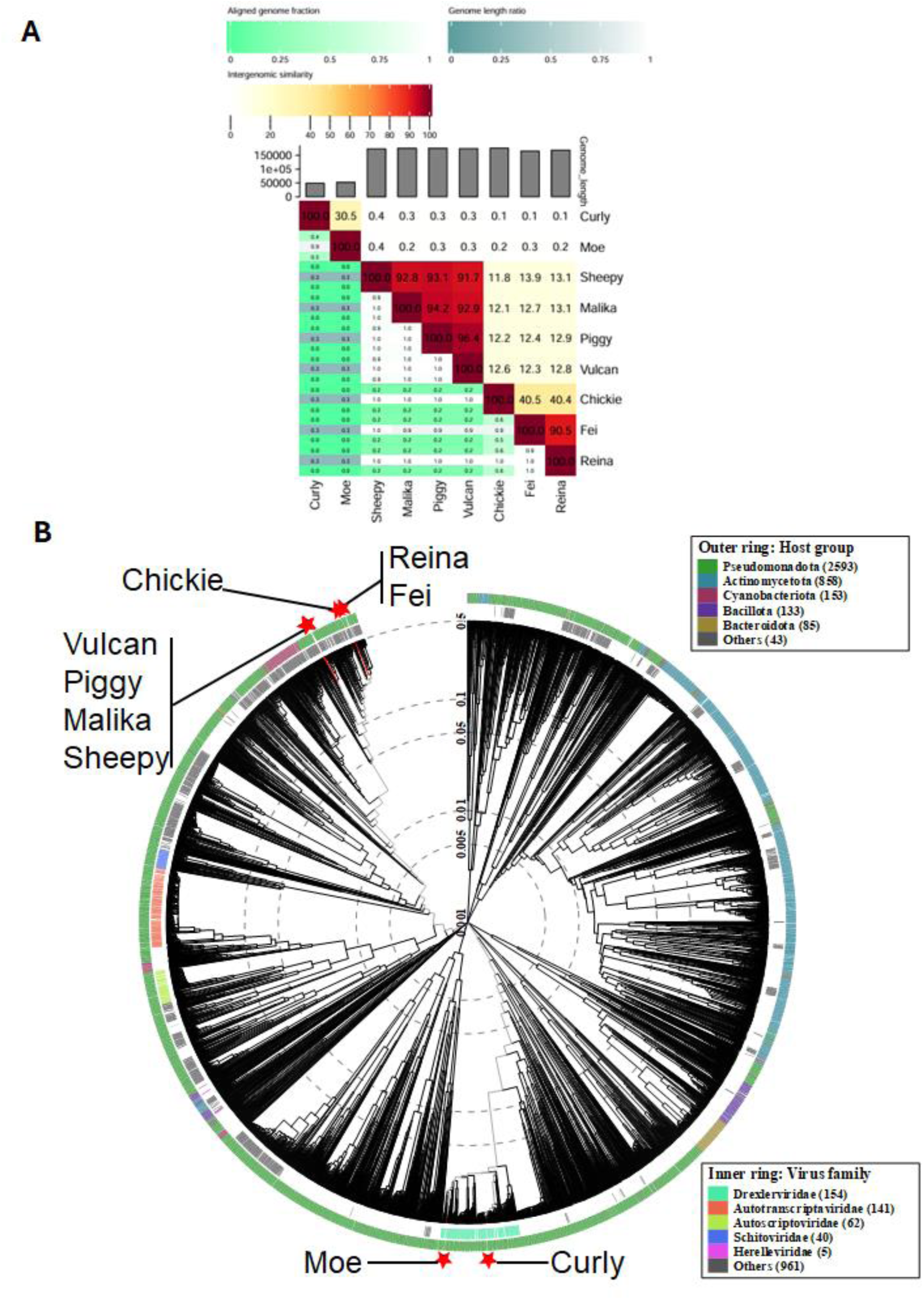
Genomic and proteomic comparison of phage isolates. **A)** Heatmap generated using VIRIDIC showing intergenomic similarity values (right half) and alignment indicators (left half and top annotation). In the right half, the color-coding shows clustering of the phage genomes based on intergenomic similarity. The more closely-related the genomes, the darker the color. White-Orange-Red color gradient was used to represent intergenomic similarity values . The numbers represent the similarity values for each genome pair, rounded to the first decimal. In the left half, three indicator values are represented for each genome pair, in the order from top to bottom: aligned fraction genome 1 (for the genome found in this row), genome length ratio (for the two genomes in this pair) and aligned fraction genome 2 (for the genome found in this column). The darker colors emphasize low values, indicating genome pairs where only a small fraction of the genome was aligned. Seagreen to white color gradient was used top and bottom indicators and Cadetblue to white color gradient was used middle indicator. The aligned genome fractions are expected to decrease with increasing the distance between the phages. Therefore, darker colors should correspond to genome pairs with low similarity values, and whiter colors to genome pairs with higher similarity values. Similarly, more closely-related viruses are expected to have similar lengths. **B)** Circular proteomic tree showing the comparison of 3924 prokaryotic virus genomes available in Virus-Host DB against the isolates in our study. Our isolates are shown as ‘red asterisks’ on the outer ring. Color rings indicate virus families (inner rings) and host groups (at a level of phylum except for Proteobacteria; outer rings). These trees are calculated by BIONJ based on genomic distance matrixes, and mid-point rooted. Branch lengths are log-scaled.

A second closely related pair was formed by Fei and Reina, which exhibited 90.5% intergenomic similarity **(Figure 1A)**. Both phages were isolated from hospital wastewater and were assigned to the genus *Jiaodavirus*. Chickie was substantially more divergent among all phages with large genome, showing approximately 40% nucleotide similarity to Fei and Reina and only approximately 12% similarity to members of the Sheepy–Malika–Piggy–Vulcan group **(Figure 1A)**. The two smaller-genome phages, Curly and Moe, were highly distinct from the remaining isolates **(Figure 1A)**. Curly, a 48,920-bp *Webervirus* isolated from horse feces, and Moe, a 52,372-bp *Peekayseptimavirus* isolated from pig feces, shared 30.5% intergenomic similarity with each other but showed only minimal nucleotide similarity to the larger-genome phages **(Figure 1A)**. Overall, the VIRIDIC analysis demonstrated substantial genomic heterogeneity within the phage collection while identifying discrete groups of closely related phages, most prominently the Sheepy–Malika–Piggy–Vulcan cluster and the Fei–Reina pair.

### Proteomic phylogeny and pairwise genome comparisons confirm distinct groups of related phages

To further assess the evolutionary relationships among the nine phage isolates, genome-wide proteomic comparisons were performed using VIPTree. The resulting circular proteomic tree placed the nine isolates at multiple positions within the reference phage phylogeny, demonstrating substantial proteomic diversity within the collection **(Figure 1B)**. The topology broadly reflected the relationships identified by VIRIDIC. Sheepy, Malika, Piggy, and Vulcan occupied a closely related region of the tree **(Figure 1B)**, consistent with their high pairwise nucleotide similarities of 91.7– 96.4%. Fei and Reina also clustered closely, in agreement with their 90.5% intergenomic similarity **(Figure 1B)**. Chickie occupied a related but distinct position from Fei and Reina, whereas the smaller-genome phages Curly and Moe were positioned in a separate region of the proteomic tree from the larger-genome isolates **(Figure 1B)**.

Local examination of the VIPTree phylogeny further showed that all nine isolates were positioned within broader proteomic neighborhoods containing previously described *Klebsiella* phages in the Virus-Host DB reference dataset **(Figure S1)**. Curly clustered among phages with genome sizes of approximately 48–50 kb, including several previously reported *Klebsiella* phages **(Figure S1A)**, while Moe occupied a neighboring but distinct cluster containing approximately 49–52-kb phages **(Figure S1B)**. Sheepy, Malika, Piggy, and Vulcan clustered together within the same large-genome proteomic lineage **(Figure S1C)**. Chickie, Fei, and Reina were located within another region of the tree containing *Klebsiella* phages as well as phages reported from other members of the Enterobacterales **(Figure S1D)**. Thus, although the nine isolates did not form a single taxa, their placement within the broader VIPTree reference phylogeny was consistent with their isolation as *K. pneumoniae*-infecting phages.

Pairwise tBLASTx genome comparisons were subsequently performed to visualize protein-level conservation among representative phage pairs. Curly and Moe, which shared only 30.5% nucleotide-level intergenomic similarity by VIRIDIC, nevertheless retained multiple homologous translated sequence blocks, although these were fragmented and distributed across the genomes rather than forming a continuous high-identity alignment **(Figure S2A)**. In contrast, comparison of the phylogenetically distant Moe and Sheepy genomes revealed very limited translated sequence similarity, with only an isolated conserved region detected **(Figure S2B)**. In contrast, extensive protein-level conservation was observed among Sheepy, Malika, Piggy, and Vulcan **(Figure S3)**. All six pairwise comparisons within this group showed dense tBLASTx connections spanning large portions of the genomes, with many homologous regions exhibiting high translated-sequence identity. The arrangement and orientation of individual homologous blocks varied between some genome pairs, but the overall abundance of high-identity matches confirmed the close proteomic relationship of these four phages predicted by both VIRIDIC and VIPTree.

A similar relationship was observed for Fei and Reina **(Figure S4A)**. Their pairwise comparison showed extensive high-identity translated sequence conservation across much of their approximately 166–168-kb genomes **(Figure S4A)**, consistent with their close position in the VIPTree phylogeny and 90.5% VIRIDIC similarity. By comparison, Chickie and Reina, which showed approximately 40% nucleotide similarity, retained a more fragmented pattern of translated sequence conservation **(Figure S4B)**. An even more distant comparison between Chickie and Sheepy showed substantially fewer conserved regions **(Figure S4C)**. Collectively, the VIRIDIC nucleotide comparisons, VIPTree proteomic phylogeny, and pairwise tBLASTx alignments consistently resolved the nine isolates into distinct genomic and proteomic groups and demonstrated progressively reduced genome-wide conservation among phages occupying more distant positions in the proteomic tree.

### Curly exhibits potent antibacterial activity against *K. pneumoniae* JJD85

The antibacterial activity of the nine bacteriophage isolates was initially evaluated by plaque assay against the clinical *K. pneumoniae* isolate JJD85. Substantial variation in plaque-forming titers was observed among the isolates **(Figure 2A)**. Curly exhibited the highest titer, reaching approximately 10^8^ PFU/mL, approximately two orders of magnitude higher than Sheepy and Piggy, which reached approximately 10^6^ PFU/mL. Reina, Fei, Vulcan, Moe, and Chickie exhibited intermediate titers of approximately 10^5^ PFU/mL, whereas Malika showed the lowest titer at approximately 10^4^ PFU/mL. Based on its superior plaque-forming activity against JJD85, Curly was selected for subsequent characterization and therapeutic evaluation.

**Figure 2.**
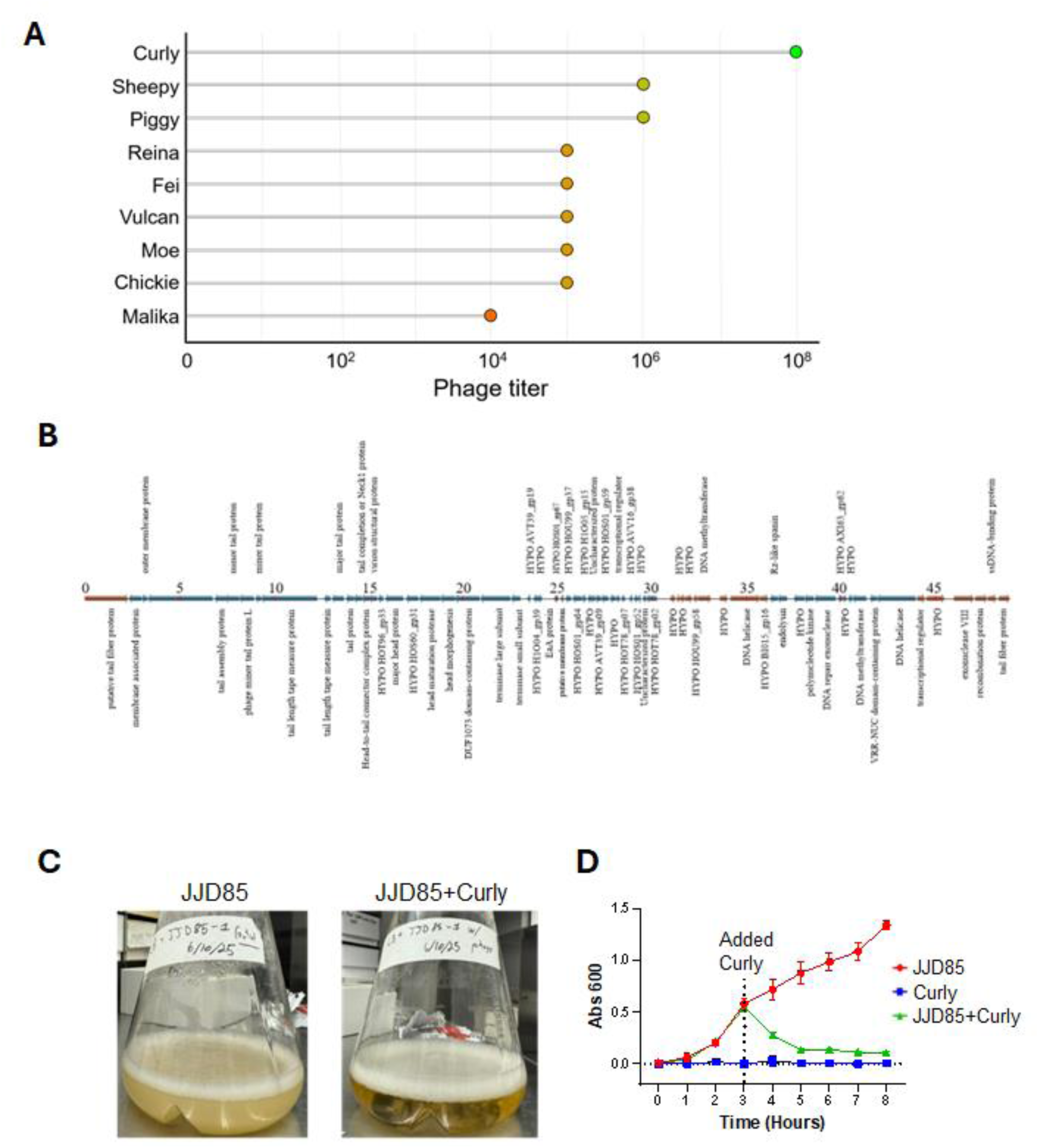
Bactericidal activity of bacteriophage isolates, and the genomic organization and antibacterial activity of phage Curly. **A)** Antibacterial efficacy of bacteriophage isolates measured by plaque assay. A lollipop plot showing plaque assay titers for individual bacteriophage isolates. Each point represents a bacteriophage isolate, and points are colored on a gradient representing low to high titer values. **B**) Predicted genome organization of bacteriophage Curly. Linear genome map of Curly showing predicted open reading frames (ORFs) across the 48,920-bp genome. Blue and brown arrows denote ORFs transcribed in opposite orientations, with arrow direction indicating the predicted direction of transcription. **C&D)** Bactericidal activity of Curly in liquid culture**. C**) Representative images of *K. pneumoniae* broth cultures after incubation in the presence or absence of Curly. **D)** Growth curves of *K. pneumoniae* measured by OD600 over time following addition of Curly during the logarithmic phase (vertical dashed line indicates the time of phage addition/infection).

Examination of the predicted protein-coding architecture of the 48.9-kb Curly genome revealed genes associated with multiple functions required for the bacteriophage replication cycle **(Figure 2B)**. The predicted structural and virion assembly repertoire included tail fiber, major and minor tail proteins, tail assembly, tail length tape-measure, head morphogenesis, and head-to-tail connector proteins, together with terminase subunits involved in genome packaging. Genes associated with DNA replication and processing included predicted DNA helicase, DNA methyltransferase, exonuclease, recombination-associated proteins, and single-stranded DNA-binding protein, in addition to several hypothetical or uncharacterized proteins. Of particular interest, the genome encodes putative tail fiber proteins, which are candidate determinants of bacterial receptor recognition and adsorption. However, the present genomic analysis does not establish whether these proteins account for the comparatively high infectivity of Curly toward JJD85.

The bactericidal activity of Curly was subsequently examined in liquid culture **(Figure 2C,D)**. Untreated JJD85 cultures showed progressive bacterial growth throughout the observation period, as demonstrated by increasing OD600. Following addition of Curly during logarithmic growth, bacterial turbidity rapidly decreased, whereas untreated JJD85 continued to grow. The OD600 of the Curly-treated culture declined markedly within the first hours following phage addition and subsequently remained near baseline, consistent with rapid bacterial lysis and sustained suppression of bacterial growth. This effect was also readily apparent macroscopically, with Curly-treated cultures exhibiting substantially greater clearing than untreated JJD85 cultures **(Figure 2C)**. Together, the plaque and liquid-culture assays identify Curly as the most active phage among the isolates tested against JJD85 and demonstrate its potent lytic activity against this clinical *K. pneumoniae* isolate.

### Curly exhibits potent antibacterial activity against *K. pneumoniae* JJD85

The antibacterial activity of the nine bacteriophage isolates was initially evaluated by plaque assay against the clinical *K. pneumoniae* isolate JJD85. Substantial variation in plaque-forming titers was observed among the isolates **(Figure 2A)**. Curly exhibited the highest titer, reaching approximately 10^8 PFU/mL, approximately two orders of magnitude higher than Sheepy and Piggy, which reached approximately 10^6 PFU/mL. Reina, Fei, Vulcan, Moe, and Chickie exhibited intermediate titers of approximately 10^5 PFU/mL, whereas Malika showed the lowest titer at approximately 10^4 PFU/mL. Based on its superior plaque-forming activity against JJD85, Curly was selected for subsequent characterization and therapeutic evaluation.

Examination of the predicted protein-coding architecture of the Curly genome revealed genes associated with multiple functions required for the bacteriophage replication cycle **(Figure 2B)**. Predicted structural and virion assembly proteins included tail fiber, major and minor tail, tail assembly, tape-measure, head morphogenesis, and head-to-tail connector proteins, together with terminase subunits involved in genome packaging. Genes associated with DNA replication and processing included DNA helicase, DNA methyltransferase, exonuclease, recombination-associated proteins, and single-stranded DNA-binding protein, along with several hypothetical or uncharacterized proteins. PhageAI classified Curly as a virulent phage with a probability of 99.85%, supporting a predominantly lytic lifestyle. The presence of putative tail fiber proteins is particularly notable because these proteins are candidate determinants of bacterial receptor recognition and adsorption; however, the present genomic analysis does not establish whether these proteins account for the comparatively high plaque-forming activity of Curly against JJD85.

The bactericidal activity of Curly was subsequently examined in liquid culture **(Figure 2C&D)**. Untreated JJD85 cultures showed progressive bacterial growth throughout the observation period, as demonstrated by increasing OD600. Following addition of Curly during logarithmic growth, bacterial turbidity rapidly decreased, whereas untreated JJD85 continued to grow. The OD600 of the Curly-treated culture declined markedly within the first hours following phage addition and subsequently remained near baseline, consistent with rapid bacterial lysis and sustained suppression of bacterial growth **(Figure 2D)**. This effect was also readily apparent macroscopically, with Curly-treated cultures exhibiting substantially greater clearing than untreated JJD85 cultures **(Figure 2C)**. Together, the plaque and liquid-culture assays identify Curly as the most active phage among the isolates tested against JJD85 and demonstrate its potent lytic activity against this clinical *K. pneumoniae* isolate.

### Curly markedly reduces *K. pneumoniae* burden in primary human monocyte-derived macrophage cultures

Because macrophages are important components of innate pulmonary defense during *K. pneumoniae* infection, we next examined whether Curly retained antibacterial activity in a host-cell environment. Primary human CD14-positive monocytes were isolated from peripheral blood of healthy donors and differentiated into monocyte-derived macrophages. Flow cytometric analysis demonstrated enrichment of the CD14-positive monocyte population from approximately 26% of total PBMCs to greater than 90% following purification **(Figure S5)**, confirming successful enrichment of the monocytes used for macrophage differentiation.

Monocyte-derived macrophages were infected with the clinical *K. pneumoniae* isolate JJD85 and subsequently treated with Curly. Bacterial burden was quantified separately in the macrophage-associated cellular fraction and the cell-free culture supernatant at 1 and 4 h after phage treatment **(Figure 3A,B)**. Curly treatment produced a marked reduction in bacterial CFU in both compartments. In the cell-associated fraction, bacterial burden was reduced by several orders of magnitude at 1 h compared with untreated cultures and remained significantly lower at 4 h, despite an increase in bacterial numbers between the two time points **(Figure 3A)**. Similarly, Curly caused a pronounced reduction in bacterial burden in the cell-free supernatant at both 1 and 4 h, whereas bacterial numbers increased substantially over time in cultures not receiving phage treatment **(Figure 3B)**.

**Figure 3.**
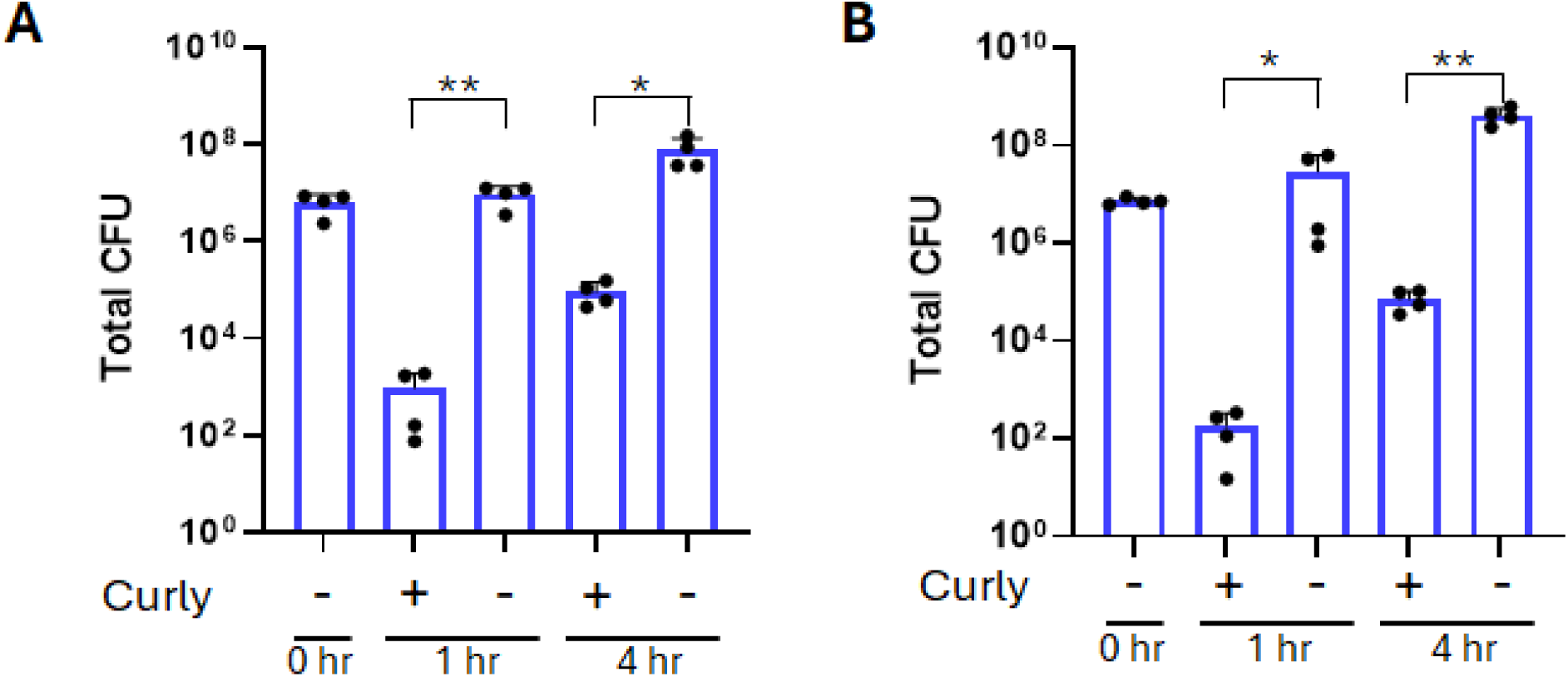
Curly reduces *Klebsiella pneumoniae* burden in primary human monocyte-derived macrophage cultures. Primary human monocyte-derived macrophages were infected with the clinical *K. pneumoniae* isolate JJD85 and cultured with or without Curly treatment. **(A)** Bacterial burden in the cell-associated fraction was quantified by colony-forming unit assay at 1 and 4 h after Curly treatment. **(B)** Bacterial burden in the cell-free culture supernatant was quantified at the corresponding time points. Data are shown for four independent donors (n = 4). Statistical significance was assessed using an unpaired t-test; *p* < 0.05 and **p** < 0.01.

These findings demonstrate that Curly maintains potent antibacterial activity in primary human macrophage cultures and restricts *K. pneumoniae* expansion in both macrophage-associated and cell-free compartments.

### Curly markedly reduces *K. pneumoniae* burden in primary human neutrophil cultures

Neutrophils are major innate immune effector cells recruited during pulmonary *K. pneumoniae* infection and contribute to bacterial clearance through phagocytosis and other antimicrobial mechanisms. We therefore investigated whether Curly retained antibacterial activity in the presence of primary human neutrophils. Neutrophils were isolated from peripheral blood and their enrichment was confirmed by flow cytometry. Following singlet selection, CD66b-positive neutrophils represented approximately 99.9% of the purified cell population, confirming successful enrichment of the cells used for subsequent infection experiments **(Figure S6)**.

Primary human neutrophils were infected with the clinical *K. pneumoniae* isolate JJD85 and subsequently cultured with or without Curly treatment. Total recoverable bacterial burden, comprising bacteria collected from both the cellular and culture supernatant components, was quantified by colony-forming unit assay at 4 and 6 h after phage treatment **(Figure 4)**. In untreated cultures, bacterial burden remained high at 4 h and increased further by 6 h. In contrast, Curly treatment resulted in a profound reduction in bacterial burden at both time points, with CFU levels remaining near 10² CFU/mL compared with approximately 10⁸ CFU/mL in untreated cultures. This corresponded to an approximately 10⁶-fold reduction in recoverable bacteria, with significant differences between Curly-treated and untreated cultures at both 4 and 6 h **(Figure 4)**.

**Figure 4.**
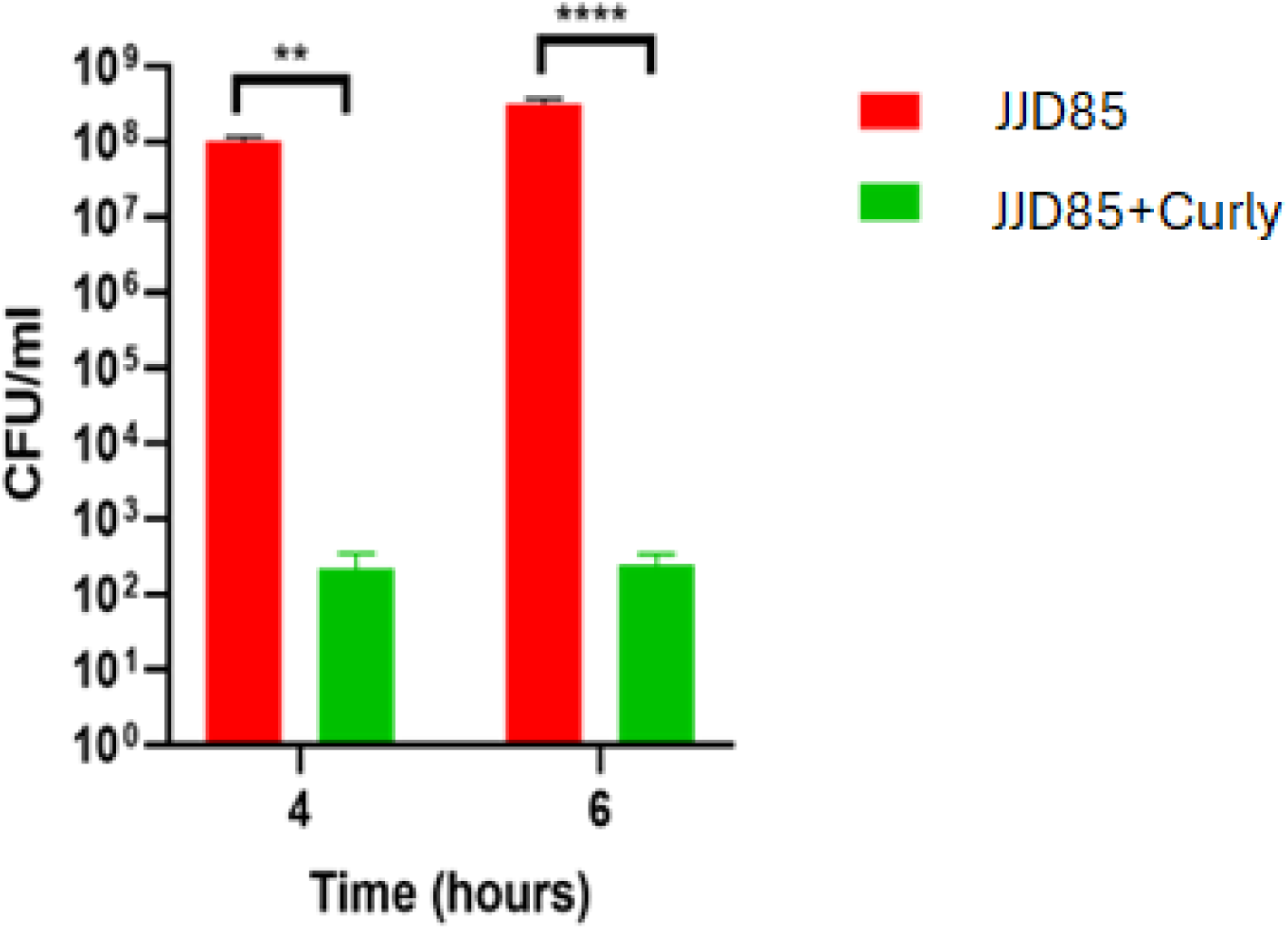
Curly reduces bacterial burden in neutrophils. Neutrophils infected with *K. pneumoniae* were treated with Curly, and total bacterial load was quantified by colony forming unit assay at 4 hour and 6-hour time points. Data are presented as mean with replicates (n = 4). Statistical significances (unpaired T-test) are denoted by asterisks (p<0.01 showing as two asterisks, and p<0.0001 showing as four asterisks).

These findings demonstrate that Curly retains potent antibacterial activity in primary human neutrophil cultures and effectively suppresses *K. pneumoniae* JJD85 despite the complex cellular environment created by a major innate immune effector population.

### TEM reveals bacteriophage infection of intracellular and extracellular K. pneumoniae

Transmission electron microscopy was used to examine the ultrastructural localization of *K. pneumoniae* and bacteriophage in infected monocyte-derived macrophage cultures. Rod-shaped bacteria were observed in both intracellular and extracellular locations **(Figure 5)**. **Figure 5A&B** show intracellular bacteria in the infected macrophages. In addition, several extracellular bacteria were closely associated with macrophage plasma membrane extensions. In **Figure 5C**, macrophage pseudopodial processes partially surrounded bacterial cells, forming phagocytic cup-like structures consistent with ongoing bacterial engulfment. Thus, the TEM images captured *K. pneumoniae* in distinct spatial relationships with macrophages, including extracellular bacteria, bacteria undergoing phagocytic uptake, and bacteria residing within macrophage-associated intracellular compartments.

**Figure 5.**
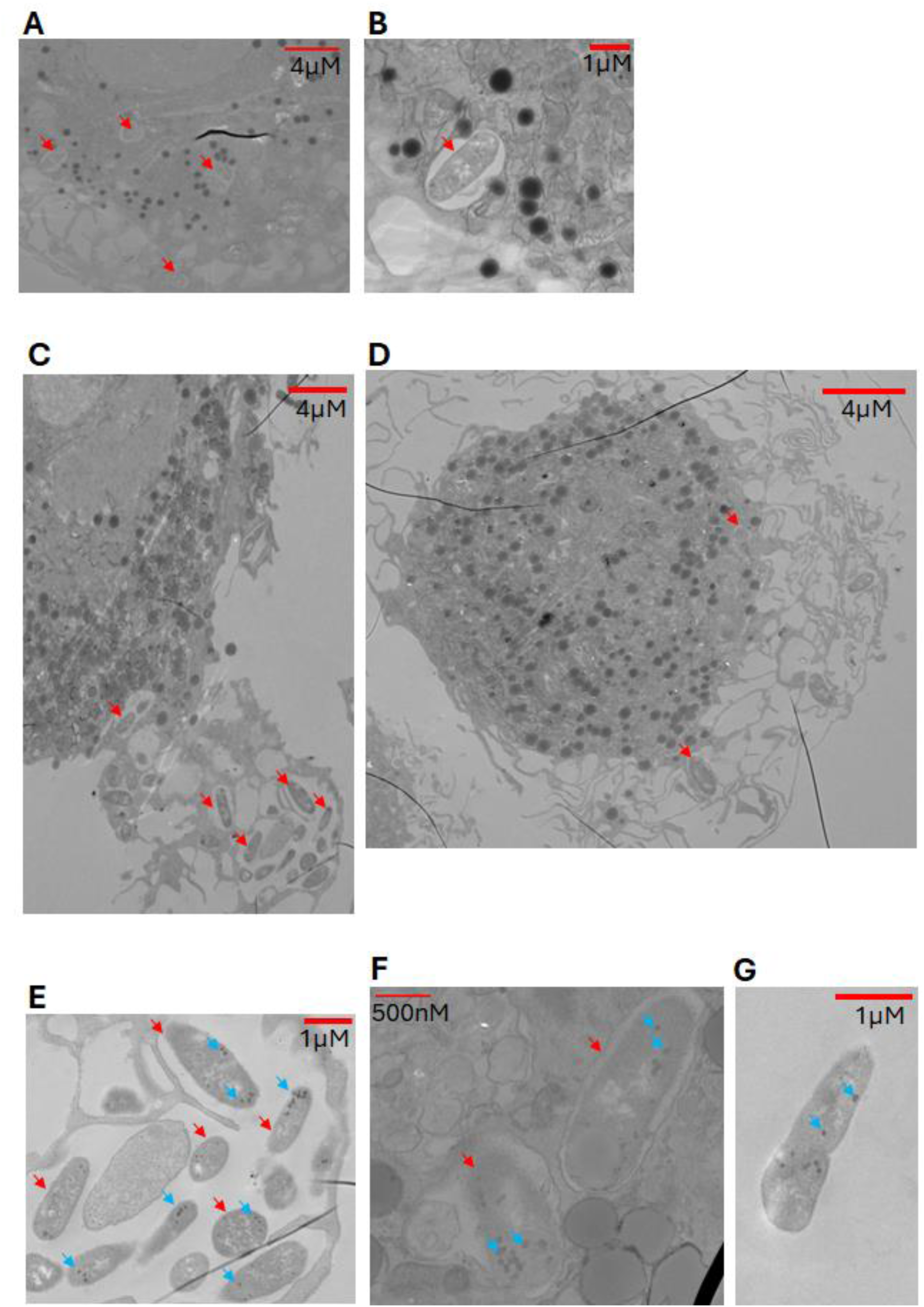
Transmission electron microscopy reveals phage-associated structures in intracellular and extracellular *Klebsiella pneumoniae*. Representative transmission electron microscopy (TEM) images of *K. pneumoniae* and bacteriophage in infected primary human monocyte-derived macrophage cultures. (A–D) Rod-shaped bacteria are observed in both extracellular and intracellular locations. In **C**, several extracellular bacteria are partially surrounded by macrophage pseudopodial membrane extensions, forming phagocytic cup-like structures consistent with ongoing bacterial engulfment. **(E–G)** Higher-magnification images show small electron-dense particles with dimensions and morphology consistent with bacteriophage heads within bacterial profiles. In phage-like particles are observed within bacteria located in **E**, macrophage pseudopodial membrane extensions, **F**, in macrophage-associated intracellular compartments, whereas **G** show phage-like particles within extracellular bacteria. Panel **G** shows an isolated extracellular bacterium containing multiple phage-like particles. Red arrows indicate bacteria and cyan arrows indicate bacteriophage heads. Scale bars: 4 µm in A, C, and D; 1 µm in B, E, and G; and 500 nm in F.

Higher-magnification TEM revealed small electron-dense particles with dimensions and morphology consistent with bacteriophage heads within bacterial profiles **(Figure 5E–G)**. Importantly, these phage-like particles were observed in bacteria in both intracellular and extracellular locations. In **Figure 5E**, phage-like particles were detected within bacterial profiles located in macrophage pseudopodial membrane extensions. In **Figure 5F**, phage-like particles were detected within bacterial profiles located in macrophage-associated intracellular compartments. Similar particles were observed within extracellular bacteria as shown in **Figure 5G** which shows an isolated extracellular bacterium containing multiple phage-like particles. Collectively, these ultrastructural observations provide morphological evidence supporting bacteriophage interaction with, and infection of, *K. pneumoniae* in both intracellular and extracellular environments.

### Curly reduces pulmonary *K. pneumoniae* burden in a murine pneumonia model

To establish a pulmonary infection model suitable for evaluating phage therapy, C57BL/6J mice were intranasally infected with the clinical *K. pneumoniae* isolate JJD85 at inoculum of 10^8^ (Low dose) or 10^9^ (high dose) CFU and monitored for disease progression. Both infection doses resulted in rapid body-weight loss during the first several days after infection **(Figure S7A)**. Mice receiving the lower inoculum subsequently showed progressive recovery, with body weight approaching that of uninfected controls by day 8. In contrast, mice infected with 10^9^ CFU showed more limited recovery and remained below baseline throughout the observation period. Consistent with these clinical changes, lung bacterial burden declined between days 2 and 4 in both groups whereas bacterial burden remained relatively stable between days 4 and 8 **(Figure S7B)**. However, in high dose mice, the bacterial burden is higher at the end of the experiment compared to low dose. Based on these findings, the higher-dose infection condition was selected for subsequent therapeutic evaluation of Curly.

To test therapeutic efficacy under conditions of sustained infection, mice were challenged intranasally with JJD85 on two consecutive days (10^9^ CFU each on both days) and subsequently treated intranasally with Curly at an MOI of 1 (low dose) or 10 (high dose) once daily for three days. All infected groups exhibited substantial early body-weight loss, and Curly treatment did not appreciably alter the initial infection-associated decline in body weight **(Figure S8A)**. Survival was also monitored throughout the experiment. Two of seven mice in the untreated infected group died by day 4, whereas all mice receiving either Curly dose survived to the end of the observation period **(Figure S8B)**.

Although this represented 100% survival in both Curly-treated groups compared with approximately 71% survival in the untreated group, the difference did not reach statistical significance.

The therapeutic effect of Curly was further assessed by quantifying pulmonary bacterial burden at the experimental endpoint. Curly treatment reduced lung CFU in a dose-dependent manner **(Figure 6)**.

**Figure 6.**
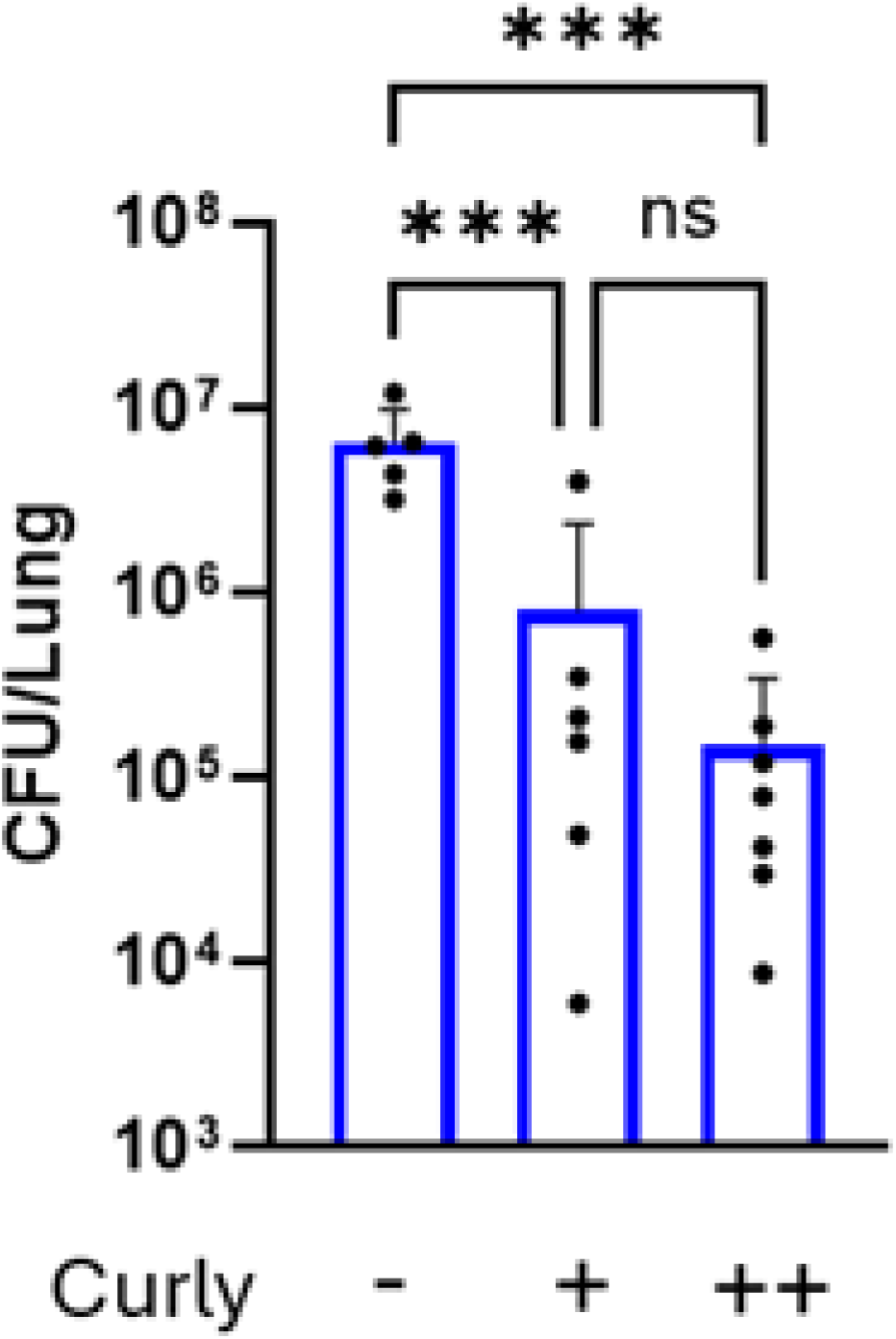
Curly reduces pulmonary *Klebsiella pneumoniae* burden in a murine pneumonia model. C57BL/6J mice were infected intranasally with the clinical *K. pneumoniae* isolate JJD85 and subsequently treated intranasally with Curly at low or high dose. Lung bacterial burden was expressed as CFU per lung. Curly treatment reduced pulmonary bacterial burden compared with untreated infected mice, with the greatest reduction observed in the high-dose treatment group. Each dot represents an individual mouse. Statistical significance was assessed using an unpaired t-test; \*\*\**p* < 0.001; ns, not significant.

Mice receiving the lower Curly dose showed an approximately 10-fold reduction in lung bacterial burden compared with untreated infected mice, whereas treatment with the higher Curly dose resulted in an approximately 100-fold reduction. Together, these findings demonstrate that Curly retains antibacterial activity in vivo and substantially reduces pulmonary *K. pneumoniae* burden in a stringent murine pneumonia model, with greater bacterial clearance observed at the higher phage dose.

Histopathological examination further demonstrated the protective effect of Curly treatment on infection-associated lung injury **(Figure 7)**. H&E-stained lung sections from uninfected control mice showed preserved pulmonary architecture, with well-defined conducting airways, open alveolar spaces, thin alveolar septa, and minimal inflammatory cell infiltration. In contrast, JJD85-infected mice exhibited extensive inflammatory cell infiltration, thickening of the alveolar architecture, and widespread parenchymal consolidation accompanied by a substantial reduction in open alveolar spaces, consistent with severe pneumonia. Curly treatment markedly attenuated these histopathological changes. Lungs from both the low- and high-dose Curly groups showed reduced inflammatory infiltration and consolidation, with greater preservation of alveolar spaces and overall pulmonary architecture compared with untreated infected mice. Residual focal inflammatory changes remained evident in phage-treated lungs, including regions surrounding conducting airways. The representative sections from the high-dose Curly group showed particularly well-preserved alveolar architecture; however, quantitative histopathological analysis would be required to establish a dose-dependent difference between the two Curly treatment groups. Collectively, these findings indicate that Curly treatment mitigates JJD85-induced pulmonary inflammation and tissue injury.

**Figure 7.**
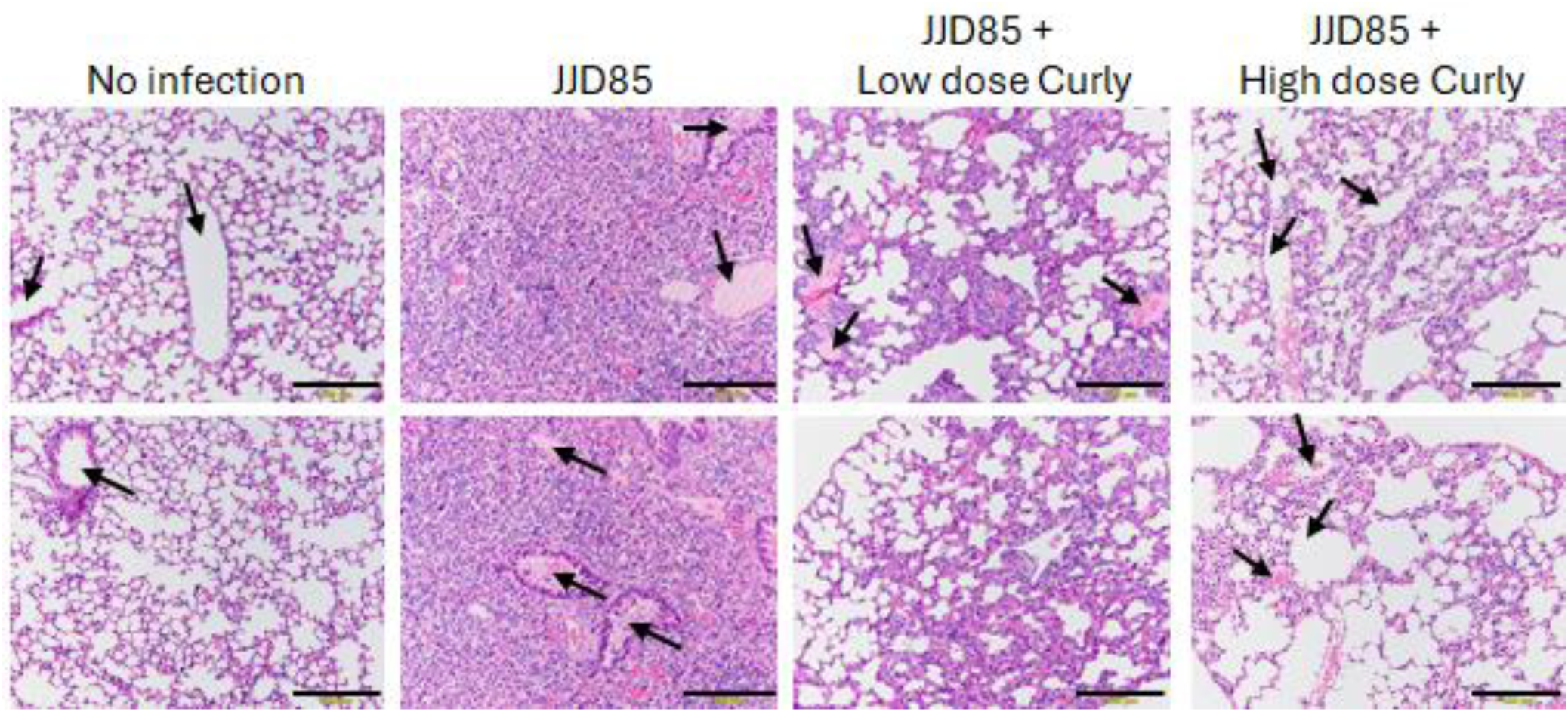
Curly treatment attenuates *Klebsiella pneumoniae*-induced lung pathology in mice. Representative hematoxylin and eosin (H&E)-stained lung sections from uninfected C57BL/6J mice, mice infected with the clinical *K. pneumoniae* isolate JJD85, and JJD85-infected mice treated with low- or high-dose Curly. Uninfected lungs show preserved pulmonary architecture with open alveolar spaces, thin alveolar septa, and minimal inflammatory cell infiltration. JJD85 infection results in extensive inflammatory infiltration, alveolar architectural disruption, and parenchymal consolidation with loss of normal air spaces. Curly treatment reduces these histopathological changes and preserves alveolar architecture compared with untreated infected mice. Black arrows indicate representative conducting airways. Scale bars, 500 µm.

## Discussion

In this study, we characterized a genetically diverse collection of Klebsiella pneumoniae-infecting bacteriophages and identified Curly as a lead candidate with potent antibacterial activity across progressively more complex experimental systems. Comparative whole-genome and proteomic analyses demonstrated substantial diversity among the nine phage isolates while identifying discrete groups of closely related phages. Functional screening against the clinical K. pneumoniae isolate JJD85 subsequently identified Curly as the isolate with the highest plaque-forming activity. Curly also rapidly suppressed bacterial growth in liquid culture, markedly reduced bacterial burden in primary human monocyte-derived macrophage and neutrophil cultures, and retained therapeutic activity in a stringent murine pneumonia model. Importantly, TEM provided ultrastructural evidence of phage-associated structures in both extracellular bacteria and bacteria located within macrophage-associated intracellular compartments. In vivo, Curly reduced pulmonary bacterial burden and attenuated infection-associated histopathological injury. Together, these findings support Curly as a promising lytic phage candidate and illustrate the value of integrating comparative genomics with functional evaluation in host-relevant experimental systems.

The comparative genomic analyses highlighted the considerable diversity that can exist among phages capable of infecting the same bacterial species. VIRIDIC resolved the nine isolates into several distinct genomic relationships, including a closely related Sheepy–Malika–Piggy–Vulcan group and a Fei–Reina pair, whereas Curly and Moe were substantially more divergent from the larger-genome isolates. VIPTree analysis broadly recapitulated these relationships at the proteomic level, and examination of the corresponding regions of the reference tree placed the isolates within broader proteomic neighborhoods containing previously characterized *Klebsiella*-infecting phages. Pairwise tBLASTx comparisons further demonstrated extensive translated-sequence conservation among phages positioned close together in the proteomic tree, whereas more phylogenetically distant pairs showed substantially less conservation. These observations are consistent with the recognized genomic heterogeneity of *Klebsiella* phages, which span multiple genomic architectures and taxonomic groups^19,28^. Such diversity may be therapeutically useful because genetically distinct phages provide a broader repertoire from which candidates with complementary host ranges or receptor specificities can ultimately be selected for phage cocktails^28–30^. This is particularly relevant for *K. pneumoniae*, in which extensive diversity of capsular polysaccharides and other surface receptors can strongly influence phage susceptibility and host range^31,32^.

An important finding was that genomic relatedness alone did not predict antibacterial performance against JJD85. Although several phages were closely related by VIRIDIC and VIPTree, their plaque-forming titers differed substantially, and Curly exhibited the highest activity despite being genomically distinct from most of the collection. This reinforces the importance of functional screening when selecting therapeutic phages^19,21^. Plaque formation and apparent phage potency can be influenced by multiple biological parameters, including receptor recognition, adsorption efficiency, latent period, replication kinetics, and burst size^20,22,33–35^. Consequently, genome size or taxonomic classification alone should not be considered a surrogate for antibacterial efficacy.

Genomic characterization nevertheless provided important information regarding the biological suitability of Curly. The 48,920-bp Curly genome contained predicted genes associated with virion structure and assembly, genome packaging, and DNA replication and processing, including tail-associated proteins, a tape-measure protein, terminase components, DNA helicase, DNA methyltransferase, recombination-associated proteins, and other predicted DNA-processing proteins. Particularly notable were predicted tail fiber proteins, because tail-associated receptor-binding structures are major determinants of bacterial recognition and host specificity ^20,35^. However, the present study does not establish that any individual tail protein accounts for Curly’s superior activity against JJD85, and direct receptor-binding or adsorption studies will be required to identify the molecular determinants of this phenotype. PhageAI independently predicted Curly to have a virulent lifestyle with 99.85% probability. Because virulent rather than temperate phages are generally preferred for therapeutic development, this prediction further supported Curly as a candidate for subsequent evaluation^10,36^. The taxonomic predictions of *Webervirus* and Drexlerviridae were supported with lower confidence and should therefore be regarded as provisional classifications rather than definitive assignments.

The marked activity of Curly in primary human immune-cell cultures is an important aspect of this study. Macrophages and neutrophils are central components of the innate response to pulmonary *K. pneumoniae* infection^24,25,37^. Macrophages participate in bacterial recognition and phagocytosis, whereas recruited neutrophils contribute to bacterial clearance through phagocytosis, reactive oxygen species, granule-associated antimicrobial mechanisms, and other effector functions^24,25,37^. *K. pneumoniae*, however, possesses multiple mechanisms that interfere with innate immune clearance, including its polysaccharide capsule, and previous studies have demonstrated that the organism can survive within macrophages by altering intracellular trafficking^26 24,38^. In our experiments, Curly markedly reduced recoverable JJD85 in both the cell-associated and cell-free fractions of monocyte-derived macrophage cultures. Curly similarly produced an approximately six-log reduction in total recoverable bacterial burden in primary human neutrophil cultures. These results indicate that Curly retains strong antibacterial activity in host-cell-associated environments that are more biologically complex than conventional broth or plaque assays. This finding is consistent with emerging evidence that phage efficacy can be strongly influenced by interactions with innate immune cells^23,39,40^.

The macrophage findings warrant particular consideration because bacteriophage access to bacteria associated with intracellular compartments remains an important mechanistic question^41,42^. The CFU experiments themselves do not establish that Curly directly entered macrophages and infected intracellular bacteria; reduction in the cell-associated fraction could potentially result from phage activity against bacteria before internalization, release and subsequent killing of bacteria from macrophages, or cooperation between phage activity and macrophage antimicrobial mechanisms.

The TEM observations, however, provide complementary morphological evidence. Bacteria were visualized extracellularly, within macrophage-associated intracellular compartments, and partially surrounded by pseudopodial extensions consistent with ongoing phagocytic uptake. At higher magnification, electron-dense structures with dimensions and morphology consistent with phage heads were observed within bacterial profiles in both extracellular and intracellular locations. These observations strengthen the possibility that Curly can interact with bacteria across different macrophage-associated compartments. Nevertheless, TEM is a static morphological method and does not by itself establish productive completion of the phage replication cycle within intracellular bacteria. Future experiments directly tracking labeled phage, quantifying intracellular infectious phage, or examining sequential stages of phage replication would be required to resolve this mechanism^41,43^. This issue is particularly relevant because interactions among phages, bacteria, and host immune cells are increasingly recognized as determinants of therapeutic outcome^23,44 41^.

The in vivo findings provide further evidence that Curly remains active in a physiologically complex infection environment. We first established a pulmonary infection model in which the higher JJD85 inoculum resulted in more sustained body-weight loss and persistent lung bacterial burden. Curly was subsequently tested in this stringent setting using repeated bacterial challenge followed by delayed phage administration. Despite these conditions, Curly reduced lung bacterial burden, with approximately a ten-fold decrease at the lower dose and an approximately 100-fold decrease at the higher dose. Previous studies have similarly demonstrated that locally administered bacteriophages can reduce *K. pneumoniae* burden in experimental pneumonia^15,16^. The dose-associated reduction observed here suggests that achieving sufficient phage exposure within the infected lung may be an important determinant of therapeutic activity^45^.

The histopathological findings complement the microbiological endpoint. Untreated JJD85 infection produced extensive inflammatory infiltration, loss of normal alveolar architecture, and pulmonary consolidation, whereas lungs from Curly-treated mice showed greater preservation of alveolar spaces and reduced inflammatory involvement. Similar reductions in pulmonary lesion severity and inflammatory infiltration following phage treatment have been reported in experimental *K. pneumoniae* pneumonia^15,46^. Thus, the therapeutic effect observed in our study was not limited to a reduction in recoverable bacteria but was accompanied by attenuation of infection-associated tissue injury. Although the representative high-dose sections appeared particularly well preserved, quantitative histopathological scoring was not performed, and a dose-dependent histological effect therefore cannot be established from these images alone. Future studies incorporating blinded semiquantitative scoring or digital measurement of inflamed and consolidated lung area would provide a more rigorous assessment of tissue protection.

Not all disease-associated outcomes improved to the same extent. Curly treatment did not appreciably prevent the early body-weight loss caused by infection. Furthermore, all Curly-treated animals survived compared with approximately 71% survival among untreated infected mice, but the difference did not reach statistical significance. Given the small group size, the study had limited ability to detect differences in survival. The dissociation between bacterial burden and short-term clinical manifestations may reflect persistence of infection-induced inflammatory responses and established tissue injury even as bacterial numbers decline. More broadly, phage efficacy in respiratory infection is determined by interactions among phage replication and pharmacokinetics, bacterial susceptibility and density, tissue delivery, and host immunity rather than bacterial lysis alone^23,40,45^.

Several limitations should therefore be considered. First, detailed therapeutic evaluation focused on a single bacterial isolate, JJD85, and one lead phage. Although the broader collection demonstrates substantial genomic diversity, the host range of Curly across genetically and capsularly diverse clinical *K. pneumoniae* isolates remains to be defined. This is particularly important because most *Klebsiella* phages exhibit restricted host ranges, with capsular diversity representing a major determinant of phage susceptibility^19,20,31^. Second, the emergence of phage-resistant JJD85 during treatment was not examined. Phage resistance is a recognized limitation of *K. pneumoniae* phage therapy, and combinations of phages with complementary receptor specificities may provide one strategy for broadening coverage and limiting bacterial escape^18,29,47^. Third, although genome annotation identified several candidate host-interaction proteins, receptor usage, adsorption kinetics, latent period, burst size, and the functions of many hypothetical proteins were not experimentally determined. These properties can substantially influence host specificity and measurable phage activity^20,22,33,34^ and could help explain why Curly displayed substantially greater activity against JJD85 than the other isolates. Fourth, the immune-cell experiments establish antibacterial activity in macrophage and neutrophil cultures but do not resolve the relative contributions of direct phage-mediated killing and immune-assisted clearance^23 40^. Finally, the murine experiments were short term and used a deliberately severe infection model; optimization of treatment timing, dose, frequency, and pulmonary delivery may produce different therapeutic outcomes ^18,45,48^.

Overall, this study demonstrates the value of combining genomic characterization with sequential functional validation when selecting phages for therapeutic development. Comparative genomics revealed extensive diversity among K. pneumoniae-infecting phages, but direct screening identified Curly as the strongest candidate against JJD85. Curly retained potent antibacterial activity in broth culture, primary human macrophage and neutrophil systems, and a murine pneumonia model, while TEM provided morphological evidence consistent with phage interaction with bacteria in both extracellular and macrophage-associated intracellular environments. The reduction in pulmonary bacterial burden together with preservation of lung architecture supports further investigation of Curly as a candidate for phage-based treatment of K. pneumoniae pneumonia. Future studies defining its host range, receptor specificity, resistance profile, pharmacokinetics, immune interactions, and efficacy in combination with other phages or antibiotics will be important for determining its translational potential.

