## Supplementary figures for "Genomic Characterization and Therapeutic Potential of the Lytic Bacteriophage Curly against *Klebsiella pneumoniae* in Human Innate Immune Cells and a Murine Pneumonia Model"

### Supplemental Materials

**Figure S1:** Portions of proteomic tree showing the immediate neighbors the tree shown in Figure 1B. Our isolates are shown as 'red asterisks' on the outer ring. Color rings indicate virus families (inner rings) and host groups (at a level of phylum except for Proteobacteria; outer rings).

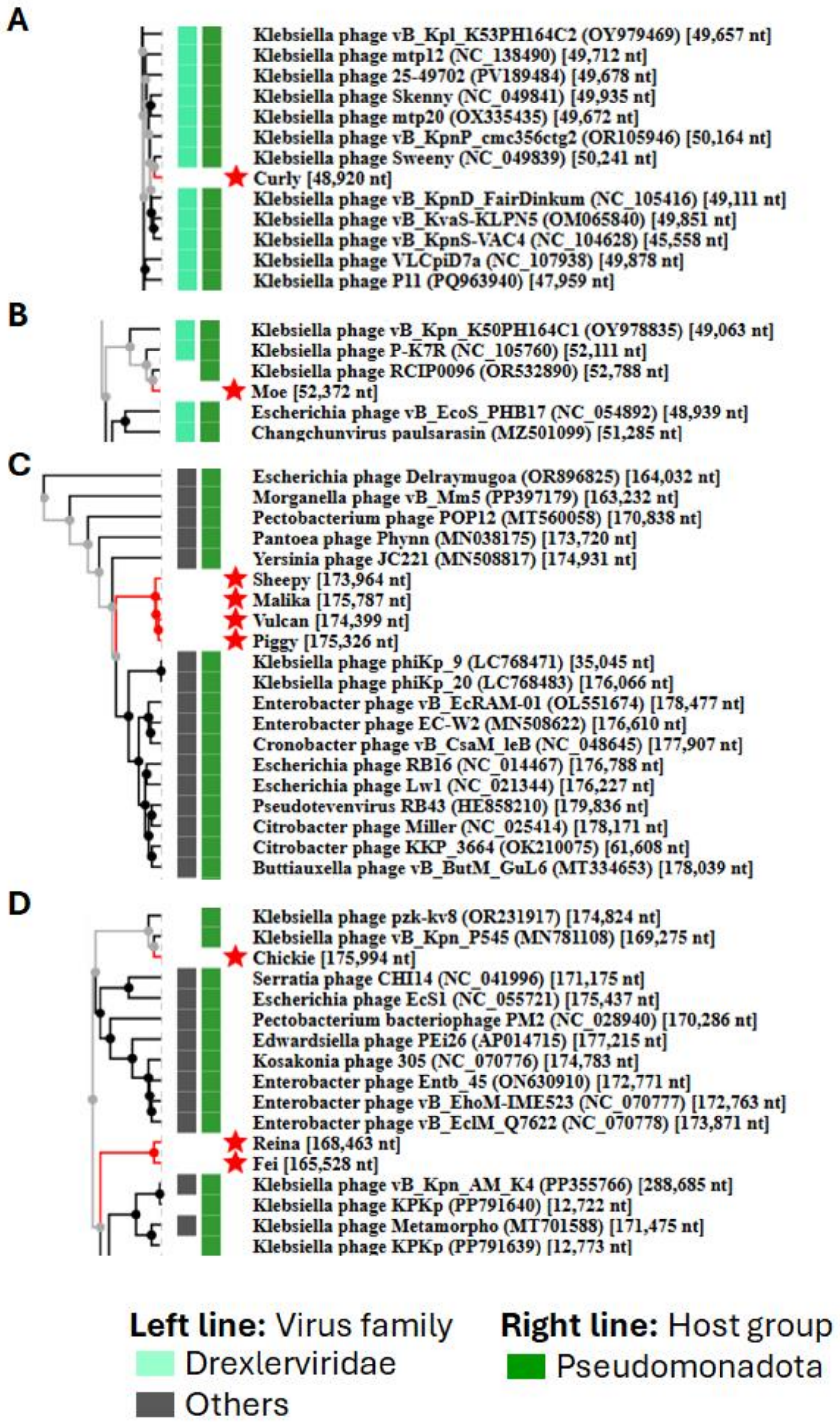

**Figure S2: Pairwise proteomic alignment view of viral genomes.** Pairwise dot plots of these genomes are also shown. **A)** Curly vs Moe, and **B)** Sheepy vs Moe. Colored lines in the alignment and the dot plots indicate tBLASTx results. Positions of each sequences are automatically adjusted (i.e., circular permuted and reverse stranded) for clear representation of collinearity between genomes.

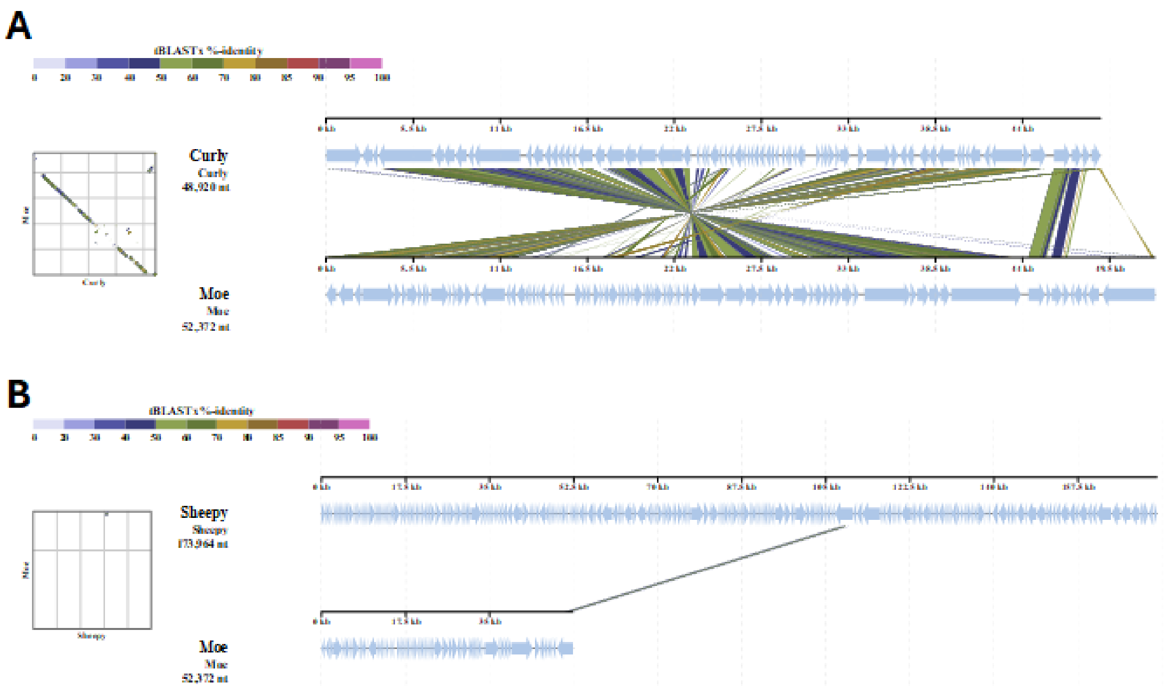

**Figure S3: Pairwise proteomic alignment view of viral genomes.** Pairwise dot plots of these genomes are also shown. **A)** Vulcan vs Piggy, **B)** Vulcan vs Malika, **C)** Vulcan vs Sheepy, **D)** Piggy vs Malika, **E)** Piggy vs Sheepy, and **F)** Malika vs Sheepy. Colored lines in the alignment and the dot plots indicate tBLASTx results. Positions of each sequence are automatically adjusted (i.e., circular permuted and reverse-stranded) for clear representation of collinearity between genomes.

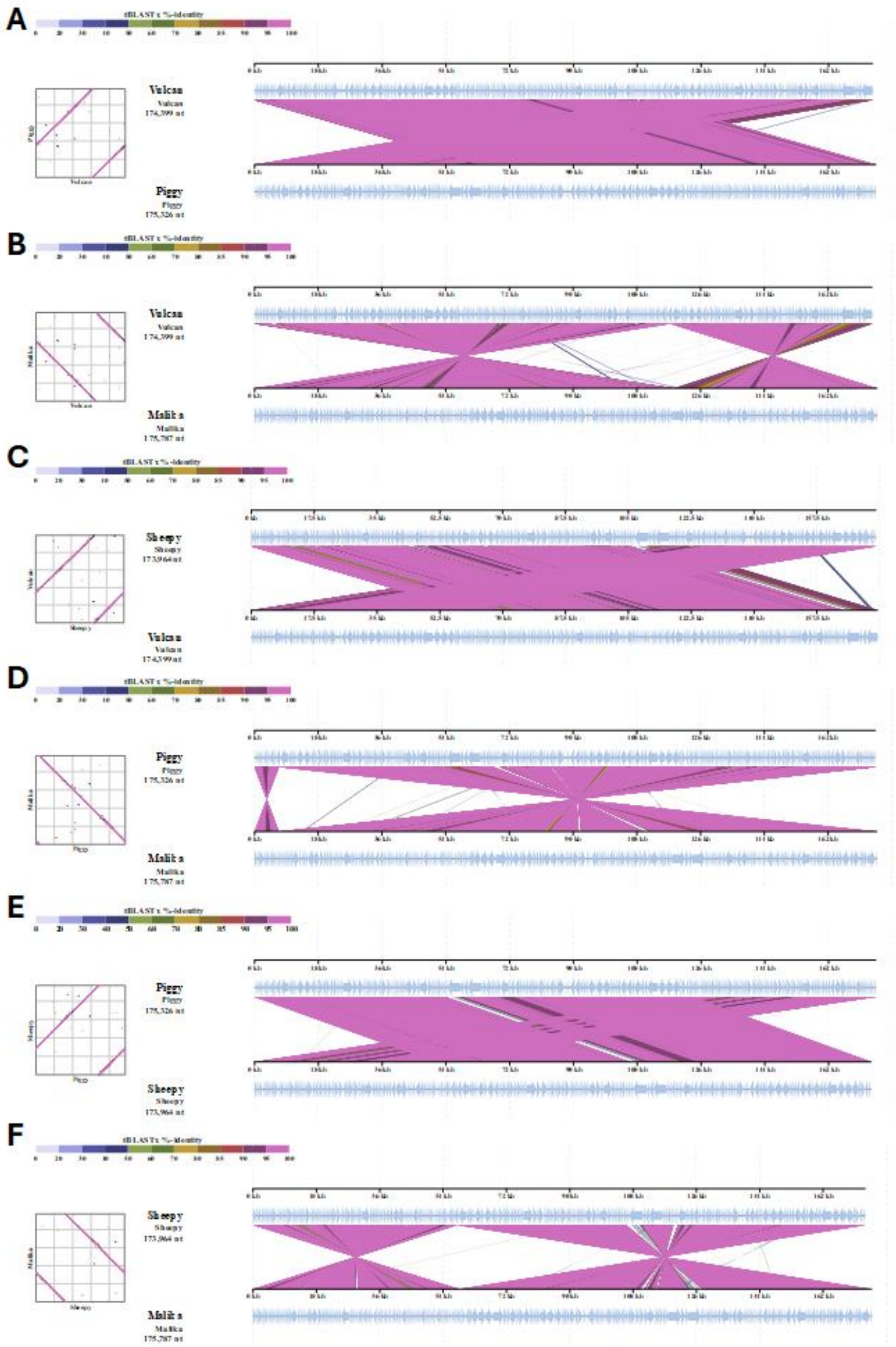

**Figure S4: Pairwise proteomic alignment view of viral genomes.** Pairwise dot plots of these genomes are also shown. **A)** Reina vs Fei, **B)** Chickie vs Reina, and **C)** Chickie vs Sheepy. Colored lines in the alignment and the dot plots indicate tBLASTx results. Positions of each sequences are automatically adjusted (i.e., circular permuted and reverse stranded) for clear representation of collinearity between genomes.

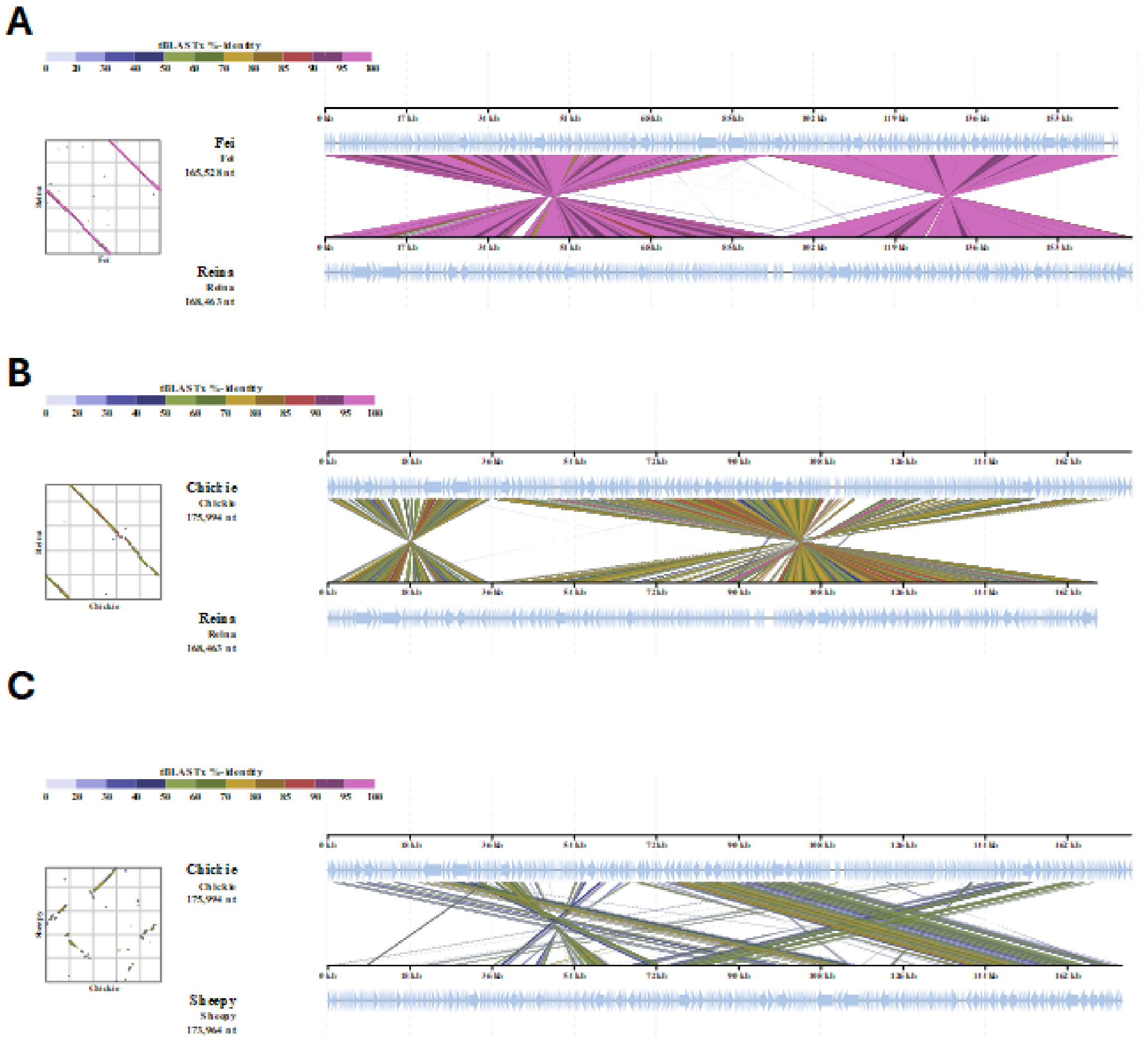

**Figure S5: Isolation and enrichment of primary human monocytes for macrophage differentiation.** Flow cytometry gating strategy for identification of CD14 positive monocytes from peripheral blood mononuclear cells. Peripheral blood mononuclear cells were first identified based on forward- and side-scatter characteristics, followed by singlet selection. CD14 positive monocytes were then gated within total PBMCs and in the enriched fraction

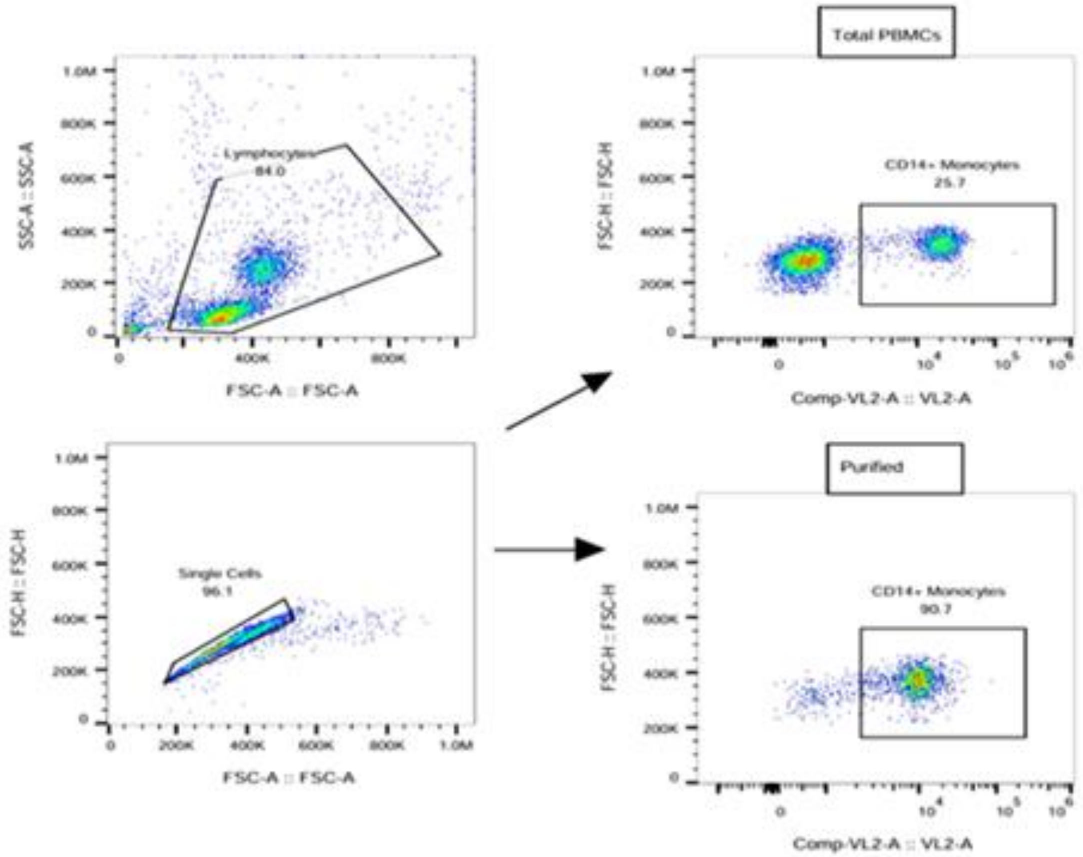

**Figure S6: Isolation and enrichment of primary human neutrophils.** Representative flow cytometry gating strategy used to assess neutrophil enrichment from peripheral blood. Neutrophils were initially identified based on forward- and side-scatter characteristics, followed by singlet selection. Following red blood cell lysis, neutrophils were identified by CD66b expression and compared before and after purification.

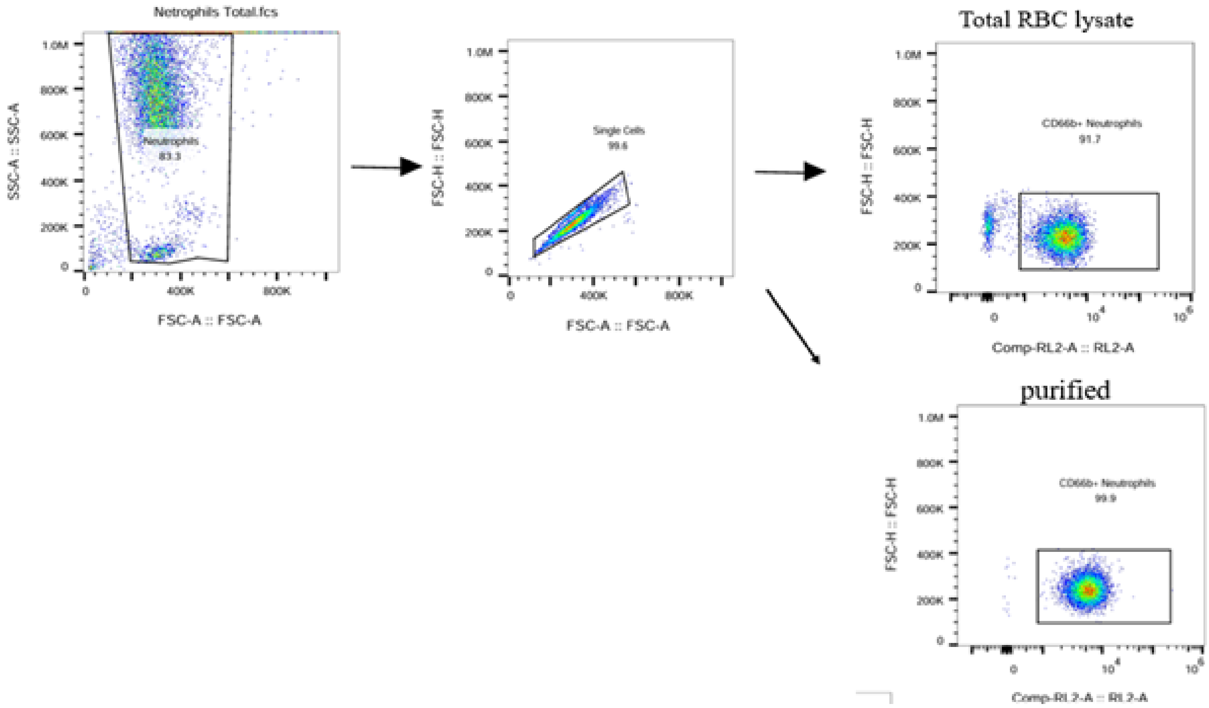

**Figure S7: Establishment of a dose-dependent murine *Klebsiella pneumoniae* pneumonia model.** C57BL/6J mice were intranasally infected with the clinical *K. pneumoniae* isolate JJD85 at a low dose (10^8 CFU) or high dose (10^9 CFU), while uninfected mice served as controls. **(A)** Body weight was monitored daily and expressed relative to baseline body weight at day 0. Both infected groups exhibited early weight loss, with greater and more sustained weight loss in the high-dose group, whereas mice receiving the low bacterial dose progressively recovered toward baseline. **(B)** Lung bacterial burden was quantified by colony-forming unit assay at days 2, 4, and 8 after infection. Each dot represents an individual mouse, and horizontal lines indicate group means. **$p < 0.01$.

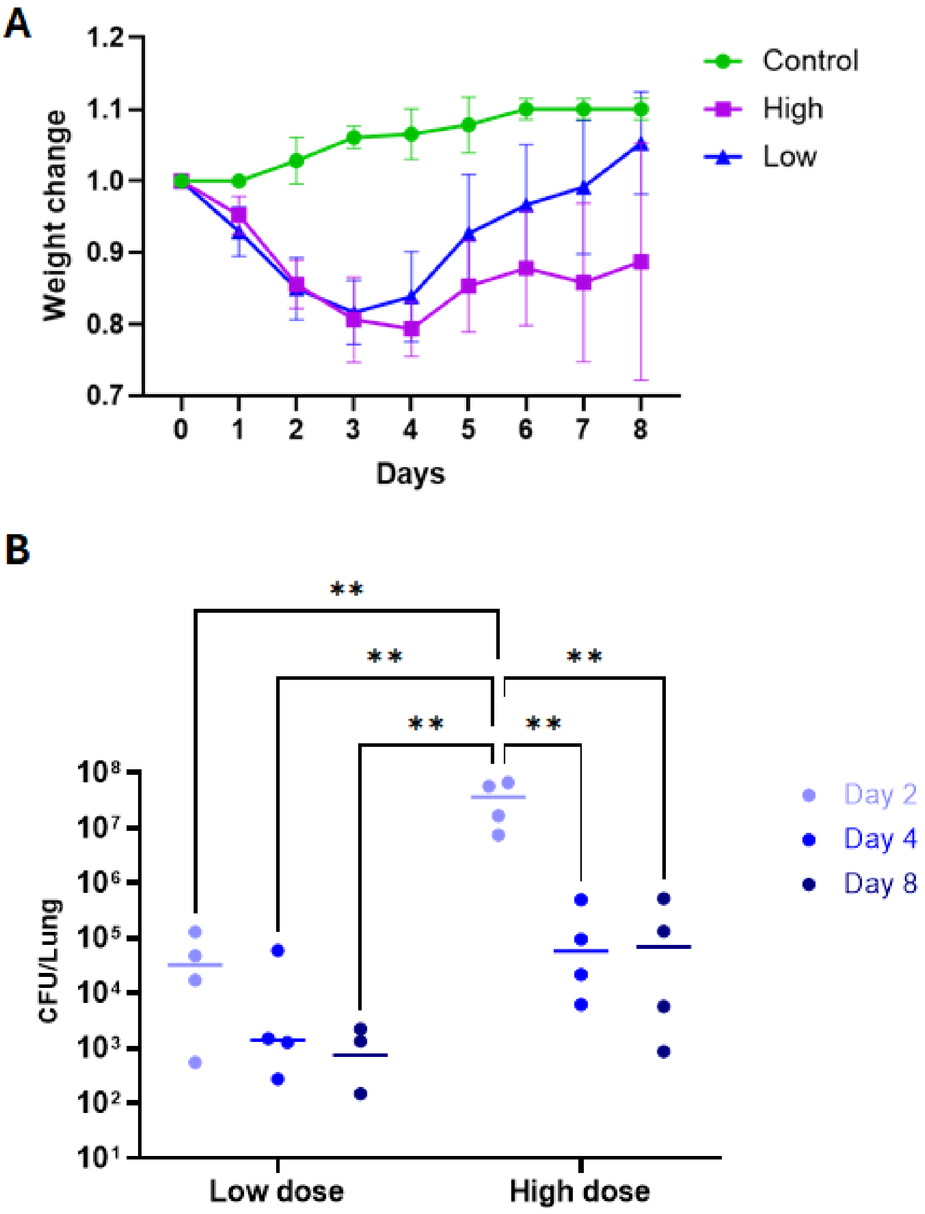

**Figure S8: Effects of Curly treatment on body weight and survival in a murine *Klebsiella pneumoniae* pneumonia model.** C57BL/6J mice were infected intranasally with the clinical *K. pneumoniae* isolate JJD85 and subsequently treated with low- or high-dose Curly. **(A)** Body weight was monitored daily and expressed relative to baseline body weight at day 0. Uninfected mice served as controls. **(B)** Kaplan–Meier survival curves for uninfected control mice, JJD85-infected mice, and JJD85-infected mice treated with low- or high-dose Curly. All mice in both Curly-treated groups survived throughout the observation period, whereas mortality occurred in the untreated infected group.

A

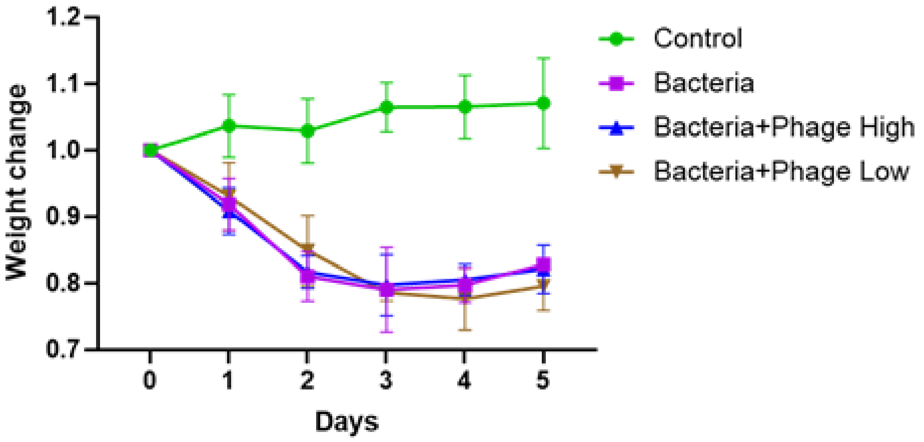


B

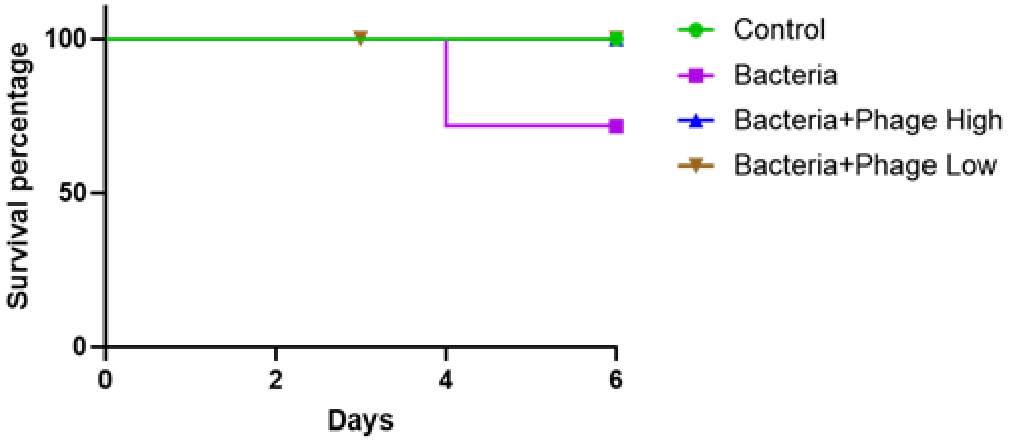
